# Deployment of non-canonical splicing in tunicate genomes is mediated by divergent U2AF function and re-patterning of snRNA m6A modification

**DOI:** 10.1101/2025.08.15.670264

**Authors:** T.C.C. Soo, A. Leon, E. Waymel, A. Barais, B. Porta, D. Chourrout, S. Henriet

**Affiliations:** Michael Sars Centre, University of Bergen, Bergen, NO-5020, Norway

**Author notes:** These authors share first authorship (T.C.C. Soo, S. Henriet).

## Abstract

In eukaryotes, the critical function of the spliceosome is to remove introns from newly transcribed pre-messenger RNA to produce functional mRNA. Spliced segments normally begin with a GT and end with an AG, and precise intron removal relies on the recognition of terminal dinucleotides by the spliceosomal RNA (snRNA) and by the U2AF heterodimer. Here, we reveal how the modification of these molecules was instrumental for shifting spliceosome specificity, enforcing splicing accuracy in genomes where 95% of introns escape the GT/AG rule.

We show that the emergence of non-canonical introns in the *Fritillaria borealis* lineage is associated with the duplication of each U2AF subunit. Subunit paralogues can be combined to form multiple U2AF complexes that involve novel protein-protein interaction rules. Binding assays on RNA and transcriptome analysis of human cells expressing U2AF subunits show that divergent paralogues can contribute to the recognition of non-canonical splice signals at the 3’ end of introns. In transfected cells, levels of N6-methyladenosine (m6A) modification on snRNA U2 and U6 are impacted by the presence of *F. borealis* U2AF2 and by AHCYL1, a specific partner of the divergent subunit U2AF2β. In striking contrast with other species, *F. borealis* has lost m6A on the U6 snRNA while U1 snRNA has gained a stable, 5’-terminal m6A. We used structure prediction to show how these snRNA modifications could play decisive roles during the recognition of non-GT/AG introns. In conclusion, we argue that spliceosome function can be profoundly impacted by gene neofunctionalization and post-transcriptional modifications, without implying major changes to conserved genetic components.

## Introduction

After transcription, the majority of eukaryotic pre-mRNA will undergo splicing, a processing step that establishes open reading frames by removing introns and by stitching exons together. This reaction is carried out by the spliceosome, a large complex composed of snRNPs *e.g.*, ribonucleoprotein particles (RNPs) assembled around a spliceosomal RNA (snRNA). Inefficient and/or inaccurate splicing may produce aberrant transcripts that cause diseases and developmental abnormalities, and one of the main challenges to spliceosome functions is to identify the exact position of introns across thousands of pre-mRNA bases. For most eukaryotic introns, the 5’ splice site (5’ss) is a GT dinucleotide and the 3’ splice site (3’ss) is an AG. These dinucleotides and their surrounding sequences are recognized by splicing factors and snRNPs (**Fig 1A**). The recognition of the intron by the U1 snRNP is often considered as the starting step of splicing. The 5’ end of U1 snRNA shows extensive complementary to the eukaryotic 5’ss consensus, which includes the 5’GT and adjacent nucleotides located in the exon and in the intron. Accordingly, mutations in the 5’ end of U1 lead to an aberrant choice of 5’ss, producing defective transcripts that can be linked to cancers^1^. The 3’ end of introns are recognized by the U2AF complex, a heterodimer composed of two highly conserved RNA-binding proteins. The small subunit U2AF1 contacts the 3’ss while the large subunit U2AF2 binds to a polypyrimidine tract (PPT) usually found immediately upstream^2,3^. The binding of U2AF to the 3’end is required for positioning the U2 snRNP on the branch point (BP) sequence. Once U1 and U2 snRNPs are bound to the intron, the U4/U6.U5 tri-snRNP is recruited and the pre-catalytic spliceosome begins to assemble.

**Figure 1:**
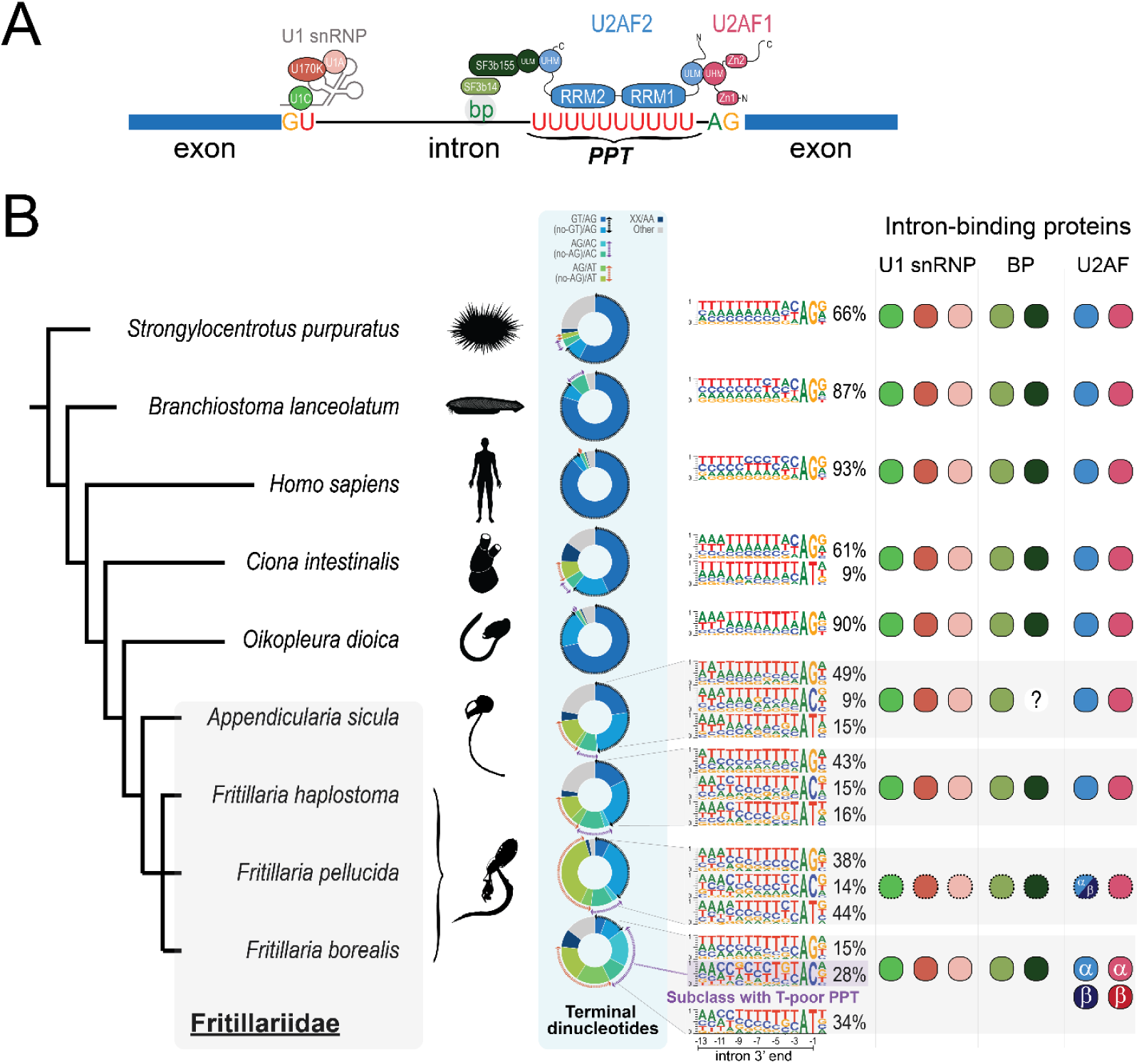
Phylogenomics of the 3’ss. **A)** Molecules involved in splice site recognition. In humans, the 5’ end of the intron is recognized by the U1 snRNP, a complex formed with snRNA, Sm proteins (not shown) and specific proteins (U1A, U1C, U1-70K). The branchpoint nucleotide (bp) is recognized by the SF3b complex (formed with subunits SF3b14 and SF3b155). The U2AF complex recognizes the 3’ end of the intron. It includes the small subunit U2AF1, that binds the 3’ss, and the large subunit U2AF2 that interacts with the PPT. Protein-protein interaction modules mediate interaction between U2AF subunits, and between U2AF and SF3b. UHM, U2AF Homology Motif; ULM, U2AF Ligand Motif. **B)** Evolution of intron classes. Species were placed on the tree based on published phylogenies^18^. The central panel shows the frequency of intron classes in different genomes, defined with their terminal dinucleotides. Statistics for *S. purpuratus*, *B. lanceolatum*, *H. sapiens* and *C. intestinalis* introns were retrieved from Minor Intron Database^61^. Logos show nucleotide composition at the 3’ end of main intron classes, along with their frequency. The right-hand panel represents gene copies identified for RNPs involved in splice site recognition. Each copy is represented by a dot, full border shows evidence gained with transcripts and genomic DNA, dashed border shows evidence from genomic DNA only. We could not find a SF3b155 gene in *A. sicula*, possibly due to incomplete genomes and transcriptomes. Greek alphabet letters represent *F. borealis* U2AF paralogues, and hybrid U2AF2 found in *F. pellucida*.

The GT/AG rule prevails in yeast and mammalian cell lines, organisms that provided key information about the mechanisms of intron recognition in the laboratory. So far, sequencing projects conducted in various species confirmed that GT/AG introns represent the major class. Because of such ubiquity, it remains difficult to assess why these dinucleotides have been selected as dominant splice site signals during eukaryotic evolution. The catalytically activated spliceosome and Group II self-catalytic introns share a conserved structure, yet these machineries process different splice sites. The so-called minor introns represent another class that corresponds to less than 1% of human introns. Minor introns are *stricto sensu* defined as being recognized by minor spliceosome (U12- type) components U11 snRNA and ZRSR2. Their respective functions are equivalent to major spliceosome (U2-type) components U1 snRNA and U2AF^4^, but they bind to different RNA sequences. While AT/AC termini represent hallmark signals for minor introns, this class of introns also includes splice sites that differ only to a little extent from the consensus of major introns. Intron removal at sites that differ substantially from the GT/AG consensus was observed in various eukaryotes, even in organisms that lack the minor spliceosome^5,6^. During evolution, splicing machineries with different selectivity have kept an active site organized around a similar architecture^7–9^. The conserved stereochemistry could define the limits of sequence evolution at intron ends, but it also implies that the spliceosome may accommodate a larger variety of splice sites than what genomes usually show.

The results gained during our last study challenged the consensus about intron recognition mechanisms, by showing that non-GT/AG introns are abundant and diverse in the larvacean tunicate *Fritillaria borealis*^10^. Non-canonical splice sites correspond to terminal inverted repeats of non-autonomous DNA transposons (“introners”) and as shown in other organisms, splicing offers a mechanism to convert gene-disrupting transpositions into introns^11^. Although other deviations have been documented in some unicellular eukaryotes^12,13^, the case of *F. borealis* is exceptional by its amplitude, since less than 10% of introns adhere to the GT/AG rule. Like other larvaceans, the U12-type spliceosome is absent in *F. borealis*^10,14^, and we could show that the U2-type spliceosome is necessary for removing both canonical and non-canonical introns. How non-canonical introns were recognized remained elusive at that time and until recently^15^, technical progress was hampered by the lack of a laboratory culture for this organism.

Apart from intron-poor organisms that tend to lose spliceosome genes, the splicing machinery has remained conserved during eukaryotic evolution^16,17^. One could assume that genome-wide modification of splice site identity would involve important changes in protein and RNA sequences that normally recognize introns, but we showed instead that the U2-type spliceosome is extensively conserved in *F. borealis* - including a strict conservation of the 5’ end of U1 snRNA. It suggests that the shift in spliceosome selectivity does not rely on major transformation of genetic components but rather on discrete, specific changes that have appeared in the *F. borealis* lineage. The experiments described here allowed us to identify when and where such modifications have been implemented during evolution, and to determine their impact on intron recognition.

## Results

### Duplication of U2AF small and large subunit genes coincides with the emergence of a new class of introns in larvaceans

Genetic resources for *F. borealis* were initially limited by specimen availability, only permitting to deploy short-read sequencing that produced genome assemblies with an important level of fragmentation^18^. We took advantage of recent developments in long-read sequencing techniques and laboratory culture to improve the genome assembly, and to establish a collection of full-length transcripts. The latest assembly features a smaller number of contigs and higher N50 (**Table S1**), which is useful for aligning full-length cDNA on the *F. borealis* genome. Based on the annotation of >30.000 introns discovered after transcript alignment, we could count only a small fraction of canonical introns (5%), while AG/AC and AG/AT were the most frequent termini seen among non-canonical introns (respectively 18 and 16%) (**Fig 1B**). To gain better insight into intron evolution in the larvacean lineage, we also sequenced DNA and RNA of two uncultured Fritillariidae species, *Fritillaria haplostoma* and *Appendicularia sicula*^15^. Even though these new genomes are still of lower quality, the long-read transcriptomes obtained with all three species allowed us to find gene orthologues and to map introns in another species, *Fritillaria pellucida*, for which no RNA-seq data is available.

Next, we compared our intron dataset to a survey of major and minor introns recently established for a variety of genomes^4^. Based on the terminal dinucleotides, we could observe a larger diversity of introns in Tunicates compared to other chordates. In agreement with earlier results^19^, intron diversity was highest in the four Fritillariidae genomes (**Fig 1B**). However, for the majority of Tunicates genome, the most frequent 3’ss corresponds to the dinucleotide AG, present in GT/AG introns and nearly canonical GX/AG introns (with X being either C, G or A) – with frequencies ranging from 43% in *F. haplostoma* up to 90% in *O. dioica*. We found a typical, uridine-rich PPT upstream the 3’AG, which is also present in other tunicate introns ending with 3’AC or 3’AT. Introns of the *F. pellucida* and *F. borealis* genomes were exceptions, since the 3’AG is less frequent than other termini (respectively 38% and 15%). Instead, the most frequent 3’ss used in these genomes is a 3’AT, found in 44% of *F. pellucida* introns and 34% of *F. borealis* introns, which is preceded by a regular PPT. In *F. borealis*, the 3’AC is the second highest 3’ss in frequency (28%), while the frequency in other species does not exceed 15% of introns. Unlike other species, the 3’AC intron subclass in *F. borealis* is characterized by a cytidine-rich PPT often interrupted by purines.

The atypical PPT of 3’AC introns in *F. borealis* resembles the PPT of minor-like introns^4^, which is at odds with the lack of U12-type spliceosome in this species. However, examining the gene complement revealed that *F. borealis* is the only species with two versions of U2AF1 and U2AF2 genes. Versions U2AF1α and U2AF2α share high sequence identity with the single copy that is found in other deuterostomes, while versions U2AF1β and U2AF2β show higher divergence (**Fig 2A**). Most differences were found in terminal arginine- and serine-rich (RS) domains, while RNA-binding domains are better conserved (**Fig 2B**). The *F. borealis* paralogues were found expressed at comparable levels in the transcriptome and *in situ*, ruling out that divergent versions correspond to pseudogenes (**FigS1**). While *F. pellucida* U2AF1 is homologous to the conserved U2AF1α, we could evidence that its U2AF2 version corresponds to a chimera between U2AF2α and U2AF2β, indicating that the common ancestor with *F. borealis* possessed two copies, that merged into one in *F. pellucida* (**Fig 2C**, **FigS2**).

**Figure 2:**
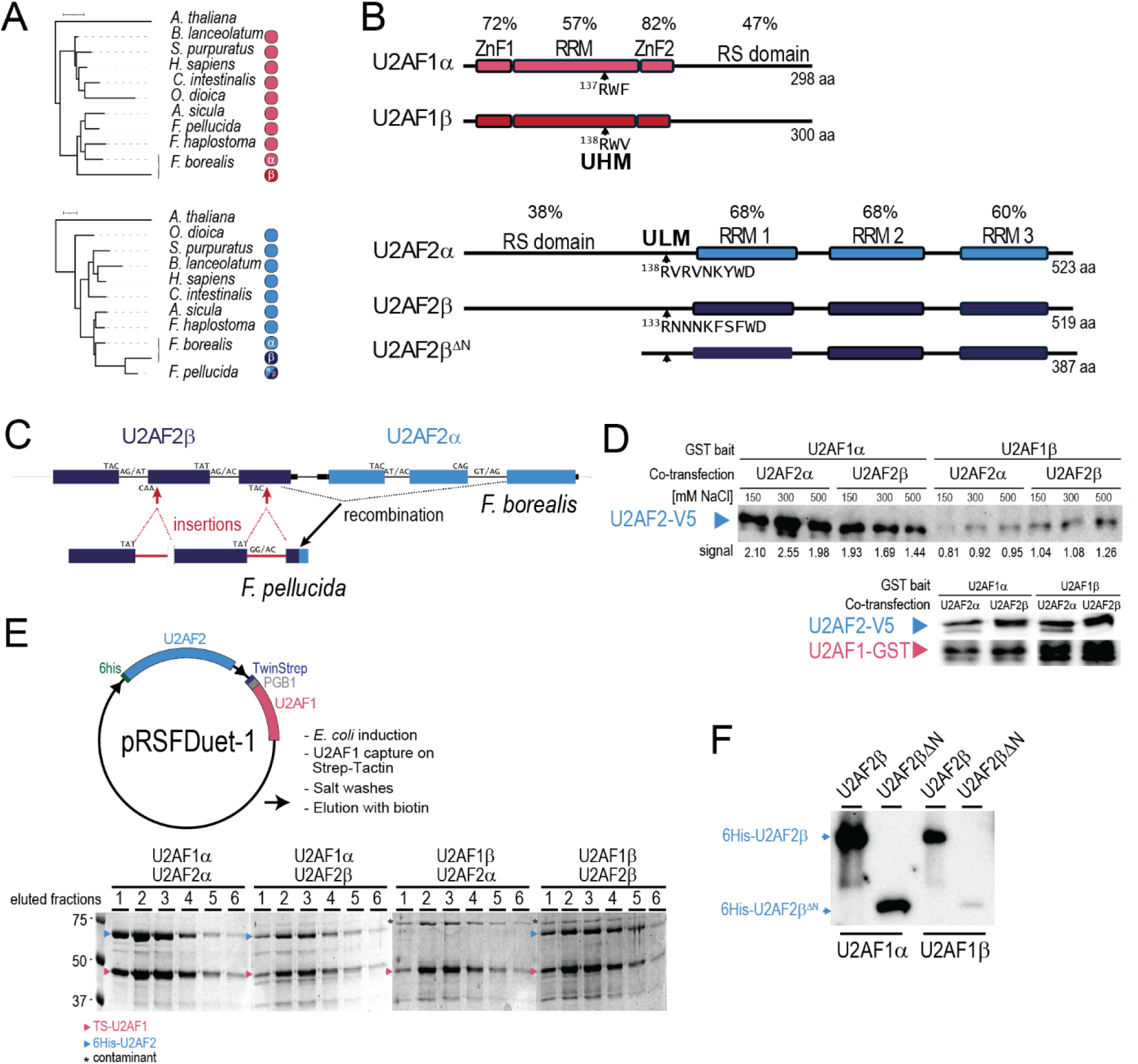
The *F. borealis* U2AF proteins. **A)** Maximum-Likelihood phylogeny of U2AF1(top) and U2AF2 (bottom) orthologues. Scales correspond to 0.1 substitution per site. **B)** Domain organization and protein-protein interaction modules in *F. borealis* paralogues. Percentage identity between paralogue domains is shown. **C)** Duplicates of U2AF2 form a gene cluster in *F. borealis*. The *F. pellucida* U2AF2 gene has features of U2AF2α and U2AF2β, showing a secondary reduction of the cluster by recombination between duplicates. Rearrangements in *F. pellucida* also include loss and gain of introns, and a reduction of the N-terminal RS domain. **D)** GST-pulldown assays in HEK293T cells. U2AF1-GST baits were co-expressed with U2AF2 tagged with a V5 epitopes. Left panel, after capturing U2AF1 on GSH-sepharose beads, associated U2AF2 was detected with an antibody against V5 (blue arrow). GST capture and washes were performed under increasing salt concentration. Right panel, expression of transfected U2AF1 (pink arrow) and U2AF2 (blue arrow) in GST-pulldown inputs. **E)** Purification of U2AF heterodimers. Left panel, representation of the construct used for tandem expression of U2AF subunits in *E. coli* and outline of the purification procedure. Right panel, Coomassie-stained SDS-PAGE showing U2AF subunits eluted from Strep-Tactin resin. Dimer formation failed only for the U2AF1β/ U2AF2α combination. **F)** U2AF1β of the RS domain abolishes dimer formation between U2AF1β and U2AF2β. Proteins eluted from Strep-Tactin resin were detected with an antibody against 6his, after testing dimer formation between U2AF1 paralogues and versions of U2AF1β.

### Divergent U2AF subunits form heterodimers with novel RNA binding properties

In principle, four different heterodimers could be formed if all the paralogues of U2AF subunits were expressed in the same cell. Sequence divergence between paralogues suggests these complexes could be associated with distinct biochemical properties, providing new functions during splicing and a division of labor. We supposed that if various U2AF possessed different affinities for RNA, they could provide a flexible way to recognize sequence diversity at the 3’ end of *F. borealis* introns, which includes the canonical combination of 3’AG preceded by a uridine-rich PPT and other combinations established with other 3’ss and the cytidine-rich PPT.

To explore this hypothesis, we first tested if different combinations of U2AF1 and U2AF2 paralogues could form protein complexes. The assembly of U2AF is mediated by interactions between the U2AF homology motif (UHM) of U2AF1 and the U2AF ligand motif (ULM) of U2AF2^20^, which we could detect in the *F. borealis* paralogues (**Fig 2B**). Using tagged U2AF1α as bait, we could recover U2AF2α and U2AF2β co-expressed in mammalian cell culture (**Fig 2D**, lanes 1-6). In comparison, the binding to U2AF2 paralogues was strongly reduced with the bait U2AF1β (**Fig 2D**, lanes 7-9). One likely reason is the replacement of the conserved F139 of the UHM by a V140 in U2AF1β, which cannot establish π−interaction with the Tryptophane present in the U2AF2 ULM^20^ (**FigS3**, **FigS4**). However, we could observe that U2AF1β interacts with U2AF2β slightly better than with U2AF2α, and that complexes formed with U2AF1β are stabilized by monovalent salts – unlike those formed with U2AF1α (**Fig 2D**, compare lanes 4-6 with lanes 10-12). Using a bacterial expression system, we could purify the two complexes formed with U2AF2α (U2AF1α/2α, U2AF1α/2β) but only a single complex formed between U2AF1β and U2AF2β (U2AF1β/2β) (**Fig 2E**). We could not isolate U2AF1β/2α, suggesting that the weak interaction detected in mammalian cell lysate is stabilized by other molecules absent in bacteria. Removing the N-terminal part of U2AF2β significantly reduced its interaction with U2AF1β (**Fig 2F**), suggesting that the loss of affinity associated with the mutation of UHM in U2AF1β can be rescued by unknown interactions established with the RS domain of U2AF2β.

We first tested how different U2AFs bind to RNA by using EMSA and oligoribonucleotide probes designed to mimic an intronic 3’ AG, preceded either by a canonical PPT (ORN.PyU) or a C-rich PPT (ORN.PyC). We tested a third probe derived from an *F. borealis* intron, where the 3’ss is replaced by an AU, preceded by a sequence rich in purines (ORN.AU) (**Fig 3A**). In agreement with the preference for uridine reported for human U2AF2^2^, the U2AF1α/2α complex interacted best with ORN.PyU, followed by ORN.AU and ORN.PyC (**Fig 3B**, **Fig 3C**). The other protein combinations, which include the complexes U2AF1α/2β and U2AF1β/2β and the single subunits U2AF1α and U2AF1β, interacted significantly better with ORN.AU than with other probes - for which levels of gelshift were almost undetectable (**Fig 3C**). It is still unclear to us whether the reduced affinity of complexes formed with U2AF2β is linked to specific substitutions present in the RNA binding domains (**FigS5**). However, when we assembled a mutant version missing the N-terminal part (U2AF2β^ΔN^) together with U2AF1α, the resulting complex U2AF1α/2β^ΔN^ could bind ORN.AU strongly, with an affinity close to the one measured for U2AF1α/2α on ORN.PyU (**Fig 3C**, **Fig 3E**).

**Figure 3:**
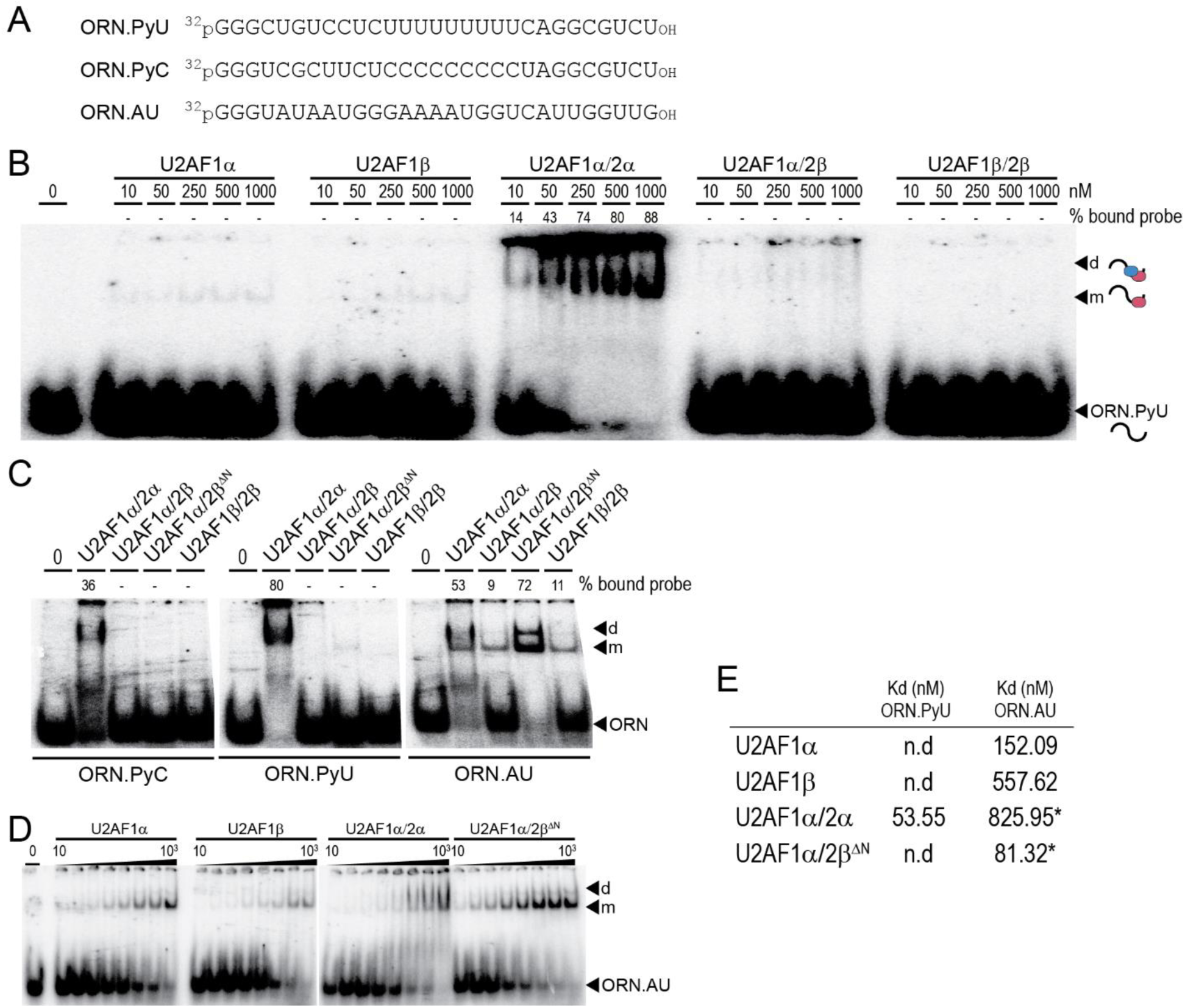
U2AF binding on RNA tested with EMSA. **A)** Representation of RNA probes used for EMSA. **B)** EMSA with ORN.PyU and purified U2AF. From left to right, experimental series were performed with U2AF1 only, or with different U2AF dimers. Bound probe measurement is reported only for values superior to 5%. Drawings on the right represent, from bottom to top: unbound probe, probe bound to one U2AF subunit and probe bound to two U2AF subunits. **C)** Comparison of U2AF binding on different probes. For each assay, 330 nM U2AF dimer was used. **D)** EMSA with ORN.AU. **E)** U2AF dissociation constants determined for ORN.PyU and ORN.AU. Values marked with a star correspond to the average of two titrations.

We predicted that sequence divergence between the conserved U2AF1 and U2AF1β could have an impact on the interaction with RNA. In the first zinc finger of U2AF1α, S38 corresponds to a highly conserved residue (S34 in human U2AF1) that is replaced by P39 U2AF1β (**FigS3**). Based on the structure of yeast U2AF1 bound to UAGGU^21^, the substitution in U2AF1β could interfere with the recognition of the position preceding the 3’AG at the end of the canonical introns (**FigS6**). We tested the binding of U2AF1 paralogues to RNA with an approach based on RNAcompete^22^. The U2AF1 paralogues were immobilized on beads, and we tested their interaction with a collection of short RNA probes derived from the 3’ ends of *F. borealis* introns (**Fig 4A**). We examined base composition among the high affinity probes, by looking first at the frequency of trinucleotides combinations with a penultimate A. Compared to U2AF1α, trinucleotides ending with AG are largely depleted in RNA recovered with U2AF1β, whereas those ending with AC are most frequent (**Fig 4B**). In U2AF1α-bound RNA, UAG triplets are more frequent than other triplets ending with AG, showing that the protein has conserved the ability to discriminate U_-3_. Dinucleotides frequencies revealed a preference for C in U2AF1β-bound RNA, and for G and U in U2AF1α-bound RNA. Similar preferences were seen for U2AF1β when comparing it to the *O. dioica* U2AF1 orthologue (**FigS7**). Based on these results and the sequence logos produced with U2AF1-bound RNA (**Fig 4C**), we propose that during splicing, the respective targets of U2AF1α and U2AF1β could be subclasses of uridine- and cytidine-rich introns.

**Figure 4:**
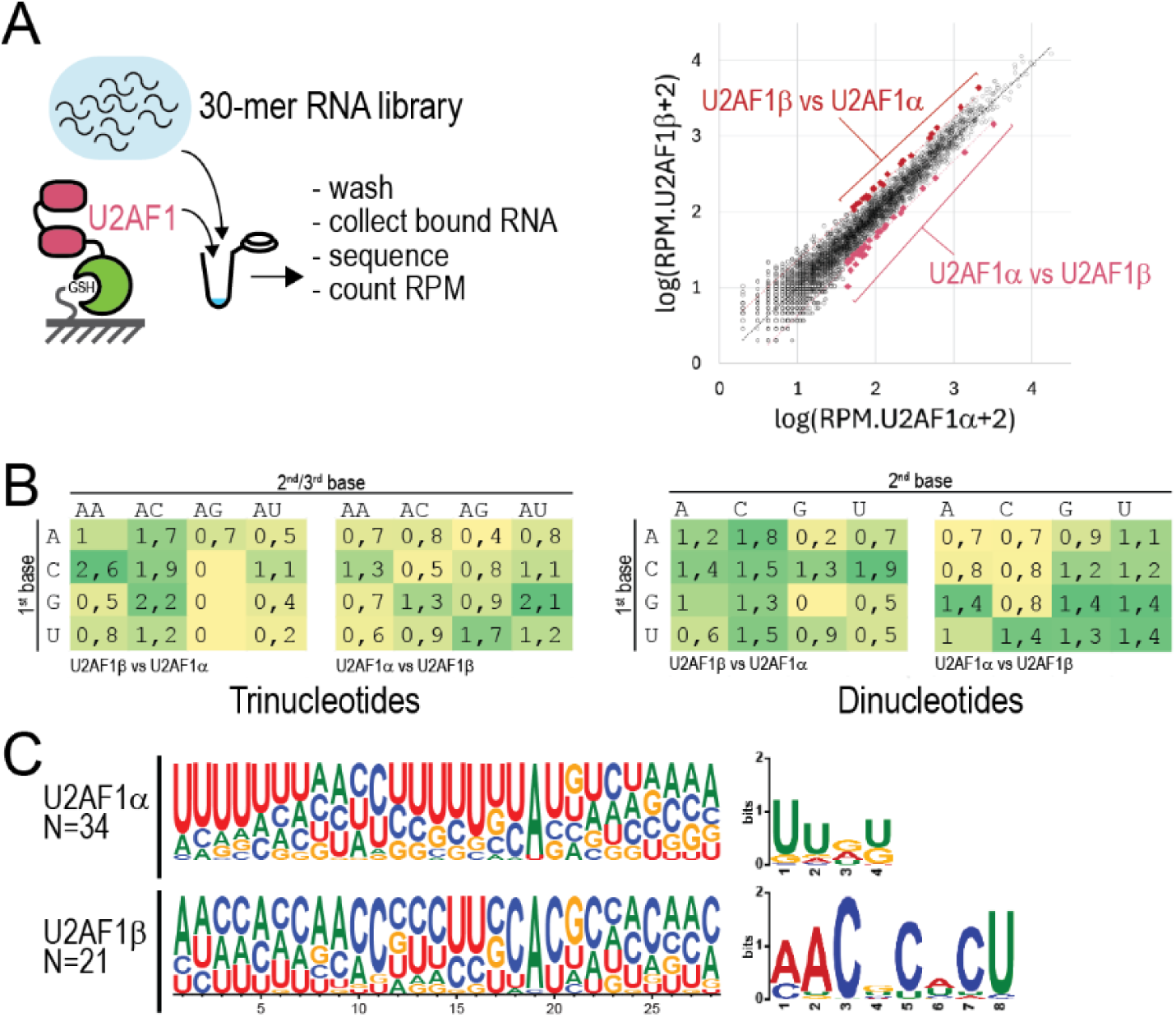
RNA-compete experiments with U2AF1. **A)** On the left, schematic drawing of approach and on the right, read count plot showing differential enrichment of library oligos in RNA recovered with U2AF1α (x-axis) and with U2AF1β (y-axis). Colored diamonds represent groups of high affinity oligos selected for sequence analysis. **B)** Abundance of 3- and 2-mer motifs in the sequence of high affinity oligos. Values correspond to the average frequency in collected oligos relative to the average frequency measured in the first library. **C)** High affinity oligos collected with U2AF1α or U2AF1β were used to establish sequence logos (left side) and compared pairwise with STREME to reveal over-represented motifs (right panel).

### U2AF2α and U2AF2β induce distinct splicing events in mammalian cells

Our results show at least three stable combinations of U2AF may exist in *F. borealis*. Depending on whether conserved or divergent versions of subunits are included, the different U2AFs are susceptible to bind to distinct types of RNA sequences. Testing U2AF function in *F. borealis* is still a challenging task, which implies developing specific approaches for genetic manipulation in this organism. Based on the good conservation of functional domains in *F. borealis* U2AF1 and U2AF2, we assumed these proteins could influence spliceosome function in a heterologous system. Expression vectors containing U2AF1 or U2AF2 paralogues were co-transfected in mammalian cell cultures, and we studied their impacts on the splicing of endogenous transcripts.

By inducing the selection of unusual splice sites, the expression of foreign splicing factors could induce the expression of transcripts carrying a premature termination codon (PTC), that are degraded by the nonsense-mediated decay pathway (NMD). To ease the detection of rare and aberrant splicing events, we first used a targeted approach to check the splicing of highly expressed genes, after a cycloheximide treatment to inhibit NMD (**Fig 5A**). The treatment allowed to increase the number of splicing isoforms and raised the detection rate of splicing events at GT/AG borders by 1.7-fold, and at other borders by 2.7- fold (**Fig 5B**). We used long-read sequencing to compare isoform expression between cells transfected with different combinations of U2AF subunits. For all samples, the expression of non-canonical isoforms (produced by the splicing of at least one GT/AG intron) remained low in comparison to canonical isoforms, but they were substantially more abundant in cells transfected with the U2AF1β/2β combination (**Fig 5C**). Each U2AF combination tested could be associated with the expression of a specific set of splicing isoforms, including those whose expression is largely reduced in other samples (**Fig 5D**).

**Figure 5:**
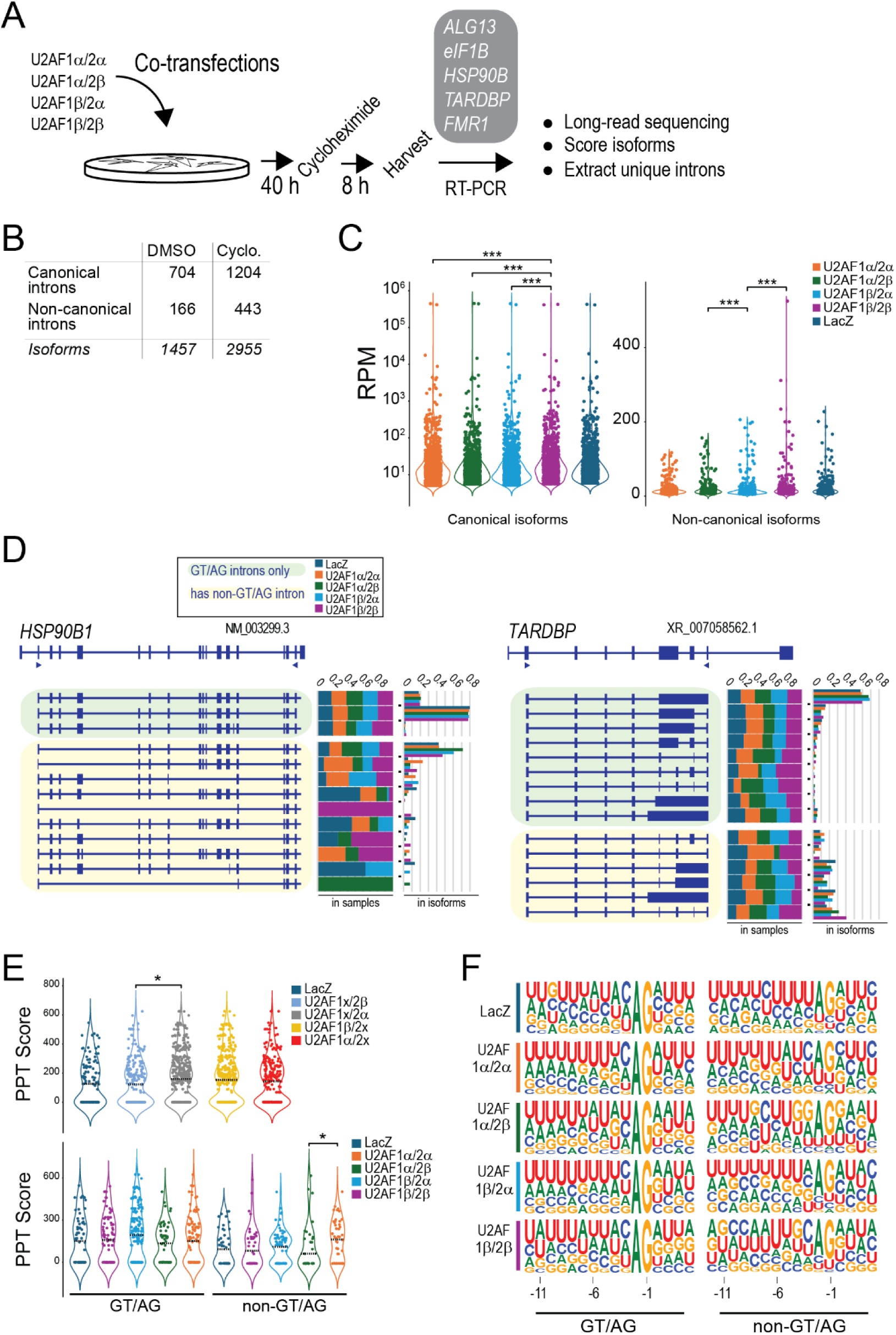
Targeted cDNA sequencing reveals different impact of U2AF paralogues on splicing. **A)** Experimental approach, showing co-transfections of U2AF subunits, NMD inhibition with cycloheximide, RT-PCR targets and isoform analysis. **B)** Table showing the total number of introns and isoforms discovered in the cDNA of either untreated cells, or cells treated with cycloheximide. **C)** Scatter plots showing read count of isoforms discovered in treated cells that are produced only by canonical splicing events (left), and for isoforms produced by the splicing of at least one non-canonical intron (right). Wilcoxon signed-rank test values are shown with stars (*, >0.05; **, >0.01; ***, >0.005). **D)** Isoform diversity in two target genes. For each gene, we represent the reference isoform on top, and the best expressed isoforms in treated cells, classified as canonical or non-canonical splicing isoforms. Arrowheads show primers used for RT-PCR. Isoform proportion is expressed either as frequency in samples, or frequency among the best expressed isoforms. **E)** Scoring PPT strength. The upper plot displays PPT scores of all introns spliced in treated cells, for control sample expressing LacZ, and for the two co-transfections made with either U2AF2β, U2AF2α, U2AF1β or U2AF1α. The lower plot displays PPT scores for distinct U2AF combinations. Dashed lines show the dataset median. **F)** Sequence logos produced with the 3’ end sequence of introns obtained from sample-specific splicing events.

Some unique isoforms were completely absent from controls cells transfected with a LacZ-expressing construct and taken together, these results show that different U2AFs induce the processing of a specific set of splice sites. For each sample, we next examined the sequence composition in unique introns, whose splicing is not detected in other samples. While no major differences appeared at the level of 3’ss, we could associate U2AF2β expression with the splicing of introns with weaker PPTs (**Fig 5E**, **Fig 5F**). Our results did not reveal a major impact of U2AF1β on splicing, perhaps due to a competition with the endogenous U2AF1, that could in principle bind to U2AF2α and U2AF2β.

We employed a regular RNA-seq approach to examine splicing globally in cells expressing U2AF2 paralogues, against a control sample that expresses a transmembrane envelope (Env)^19^. U2AF2α has a larger impact on gene expression compared to U2AF2β (**Fig 6A**).

**Figure 6:**
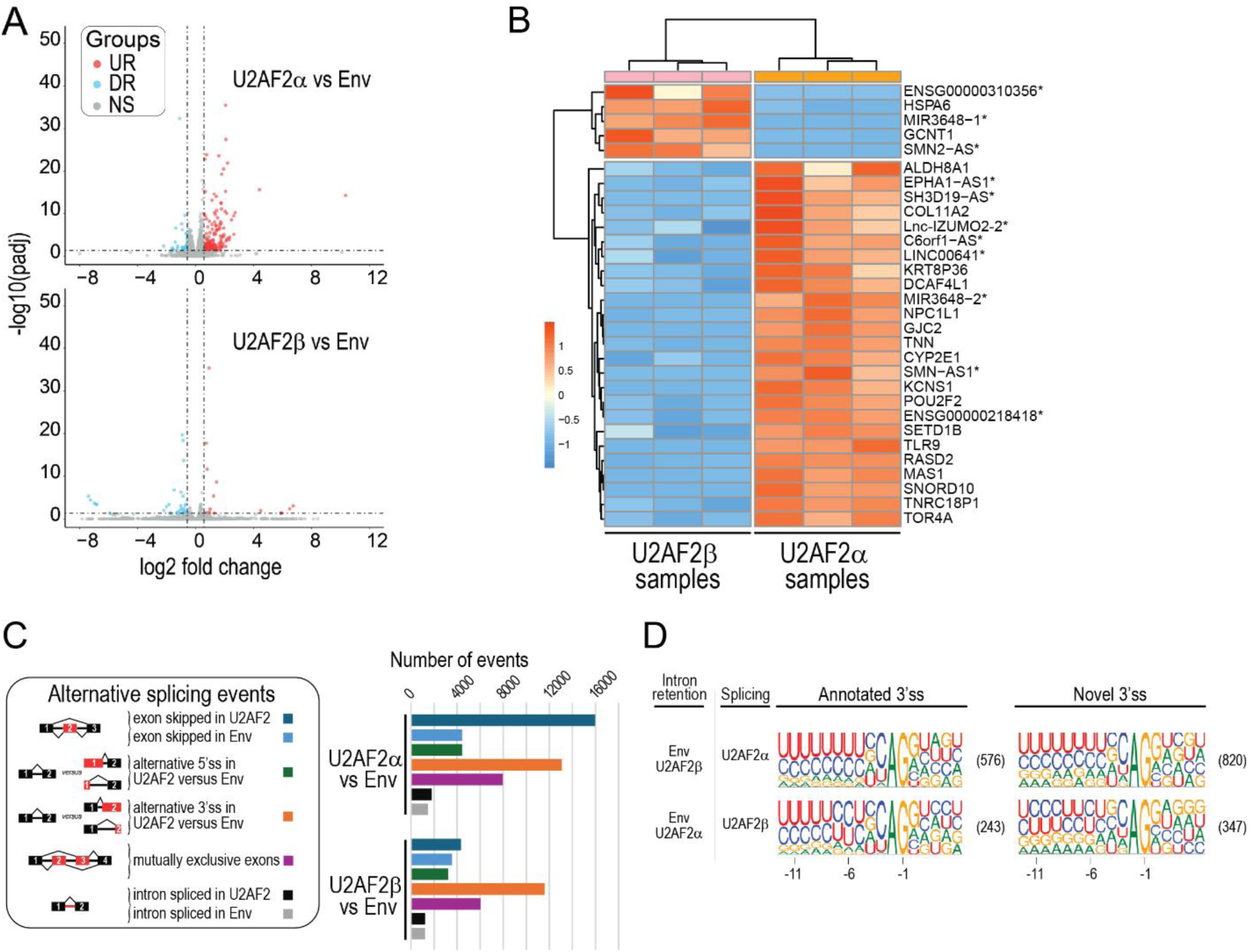
Transcriptome analysis of cells expressing *F. borealis* U2AF2. **A)** Volcano plots showing DEGs found by comparing the transcriptome of cells transfected with either U2AF2 paralogues, or control cells expressing an envelope protein. UR, upregulated genes; DR, down-regulated genes; NS, non-significant change. **B)** Top 30 DEGs between cells transfected with U2AF2α and cells transfected with U2AF2β. Stars show non-protein coding genes. **C)** AS events detected with rMATS, classified by type of events. The events include annotated and novel splice sites. **D)** Expression of U2AF2 paralogues is associated with the processing of distinct 3’ss. Sequence logos were produced with 3’ss from introns spliced specifically in cells transfected with U2AF2α or U2AF2β.

However, GO terms associated with differentially expressed genes (DEGs) primarily identified functional categories linked to protein homeostasis, suggesting effects on gene expression were provoked by the over-expression of transfected genes rather than a specific impact on transcription programs. More than a third of the top DEGs between U2AF2α and U2AF2β correspond to non-coding genes (**Fig 6B**). We interpreted this result as a possible consequence of novel splicing events induced by *F. borealis* U2AF2, leading to the degradation of numerous protein-coding transcripts by NMD and to an enrichment in non-coding RNA. Intriguingly, we saw antagonistic effects on the expression of antisense lncRNA transcribed from SMN1 and SMN2 genes, with SMN-AS1 overexpressed in U2AF2α-transfected cells and SMN2-AS overexpressed in U2AF2β-transfected cells. It is still unclear if these differences reflect the effect of U2AF2 paralogues on the processing of lncRNA or on the expression of SMN1 and SMN2 genes^23^.

After mapping reads, rMATS revealed 42931 alternative splicing (AS) events associated with U2AF2α expression, and 26373 AS events associated with U2AF2β expression. Exon skipping is largely responsible for the excess of events detected with U2AF2α, and it is the most frequent class of AS detected in this sample (15969) (**Fig 6C**). Given that U2AF2α and human U2AF2 may share similar preferences for RNA, we suppose that the presence of U2AF2α can contribute to the exon skipping activity already shown for human U2AF. The binding of U2AF to intronic sites was indeed shown to interfere with the selection of 3’ss located immediately downstream^24^. While lower concentrations of U2AF may leave these intronic sites vacant, hybrid U2AF formed with U2AF2α in transfected cells could bind these sites, leading to the strong increase in exon skipping events. Hybrid U2AF that includes *F. borealis* U2AF2α or U2AF2β and endogenous U2AF1 are likely to form in transfected cells, based on our protein interaction assays. However, the lower number of registered events suggest U2AF2β is largely unable to induce exon skipping with the same mechanism – possibly due to different affinity for RNA. We used the set of AS events identified with rMATS to check for sequence differences around the 3’ss. When examining splice sites that are specifically processed in one U2AF2 paralogue transfection but not with the other paralogue or the Env control, the only AS class showing substantial variation was intron retention. Here, splicing induced by U2AF2α is associated with uridine-rich PPT before the 3’ss (**Fig 6D**), while cytidines are more frequent in the PPT of introns spliced under U2AF2β expression. Consistent with results gained with the targeted approach, these observations show that U2AF2α or U2AF2β induce the splicing of distinct sets of introns in mammalian cells, which can be distinguished based on their PPT.

### U2AF paralogues and AHCYL1 regulate N^6^-methyladenosine levels in snRNA

Our experiments revealed an unexpected function of the RS domain of U2AF2β by showing it interferes with the binding of U2AF on RNA. This region of the protein also mediates dimerization with U2AF1β while in humans, the RS domain of U2AF2 is needed for binding to Prp19 and for transcription-coupled splicing ^25^. We supposed that the discovery of proteins bound by U2AF2α and U2AF2β could provide novel indications about their function beyond 3’ss recognition, and we used a co-immunoprecipitation assay to recover the proteins that interact with U2AF2α or U2AF2β in mammalian cells (**Fig 7A**).

**Figure 7:**
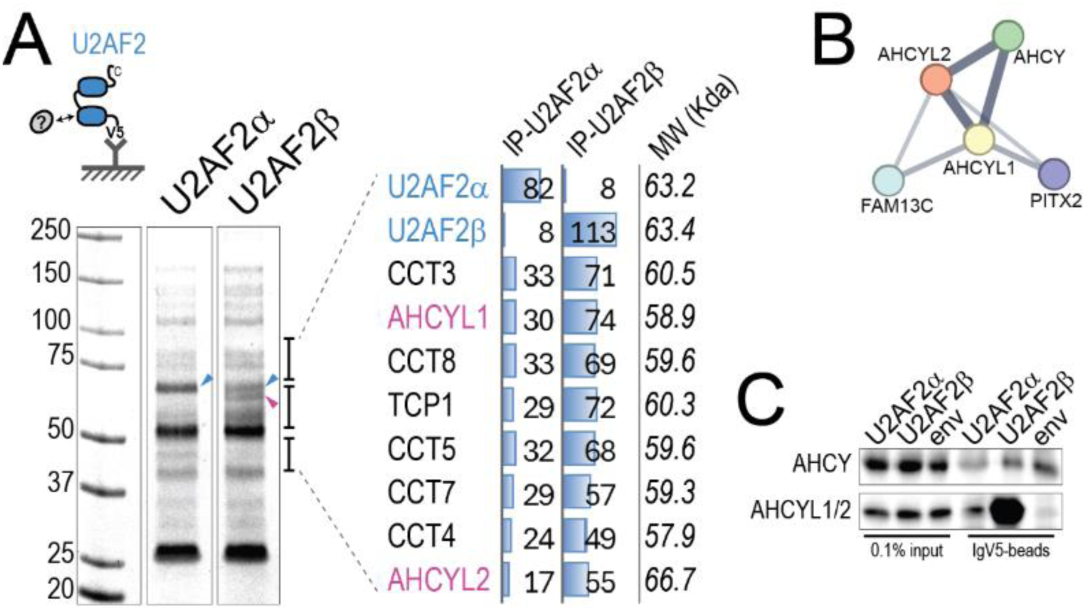
The U2AF2β-AHCYL1 interaction. **A)** Co-immunoprecipitation experiment. We expressed U2AF2 paralogues with a V5 tag in mammalian cells, and after immobilizing protein complexes on antibody-coupled beads, we examined the associated proteins on Coomassie-stained gel. Brackets show the areas analyzed with mass spectrometry. On the right-hand side, the most abundant proteins found in the analysis are listed with the number of corresponding peptides detected, and their predicted size. Blue and Pink arrows show bands corresponding to U2AF2 baits and AHCYL proteins, respectively. Note that AHCYL1 and AHCYL2 share 91.8% identity over 80% of their protein sequence. **B)** AHCYL protein interaction network retrieved from the STRING database^62^. Lines show physical interactions, with thickness proportional to data support. **C)** Validation of the interaction between U2AF2 and AHCY-like proteins. The western blot shows the result of a co-immunoprecipitation experiment using U2AF2-V5 as bait. Lanes 1-3: cell extracts expressing tagged baits, either U2AF2 paralogues, or *env* protein control. Lanes 4-6: material bound on beads after immunoprecipitation. Samples were tested with an AHCY antibody (top) or an antibody recognizing both AHCYL1 and AHCYL2 (bottom).

Mass spectrometry (MS) revealed Adenosine Homocysteinase-like proteins (AHCYL, also known as IRBIT) as a specific partner of U2AF2β. The proteins AHCYL1 and AHCYL2 (also known as SAHH2 and SAHH3, respectively) share strong sequence identity, and they are homologues of Adenosine Homocysteinase (AHCY, also known as SAHH). Diagnostic peptides identified with MS suggest AHCYL1 as the primary partner of U2AF2β. AHCYL1 and AHCYL2 have been reported to interact with AHCY (**Fig 7B**), and some of the peptides found in U2AF2β co-IP are also conserved between AHCYL proteins and AHCY. Using specific antibodies, we could confirm that AHCYL1/2 is a protein partner of U2AF2β that is efficiently recovered with co-IP (**Fig 7C**).

The enzymatic activity of AHCY contributes to stabilizing cellular pools of SAM, the substrate of methyltransferases. AHCYL1 is considered catalytically inactive and while it could interfere with AHCY function, the impact on histone methylation is still the subject of debate^26^. Interestingly, AHCYL1 and AHCYL2 bind to ZCCHC4, the methyltransferase responsible for N^6^-methyladenosine (m6A) modification of 28S rRNA, but their impact on base modification is not documented^27^. Recent evidence studies have shown that m6A can regulate splicing^28,29^ and based on our co-IP results, we decided to check the effect of AHCYL1/2 and *F. borealis* U2AF2 expression on the deposition of m6A on snRNA. Internal m6A is frequent in mammalian U2 and U6 snRNA, while for other snRNA m6A maps to the transcription start nucleotide and is kept at a low abundance by the FTO demethylase^30^.

Knocking out the expression of mammalian AHCYL1 with CRISPR/cas9 provoked a 1.5-fold increase of m6A signals on U2 and U6 snRNA (**Fig 8A, FigS8A, SI**), showing that in normal conditions AHCYL1 represses the modification of these target adenosines. For both *F. borealis* U2AF2 paralogues, expression in mammalian cells was associated with reduced m6A levels in U6, while U2AF2α specifically reduced the m6A levels in U2 (**Fig 8B, FigS8B, SI**). Although we could reveal only modest effects, our results show that AHCYL1 and U2AF2 can regulate the m6A modification of internal adenosines, opening for a new mechanism that is distinct from the FTO-mediated control of N^6^ methylation on the 5’-terminal adenine.

**Figure 8:**
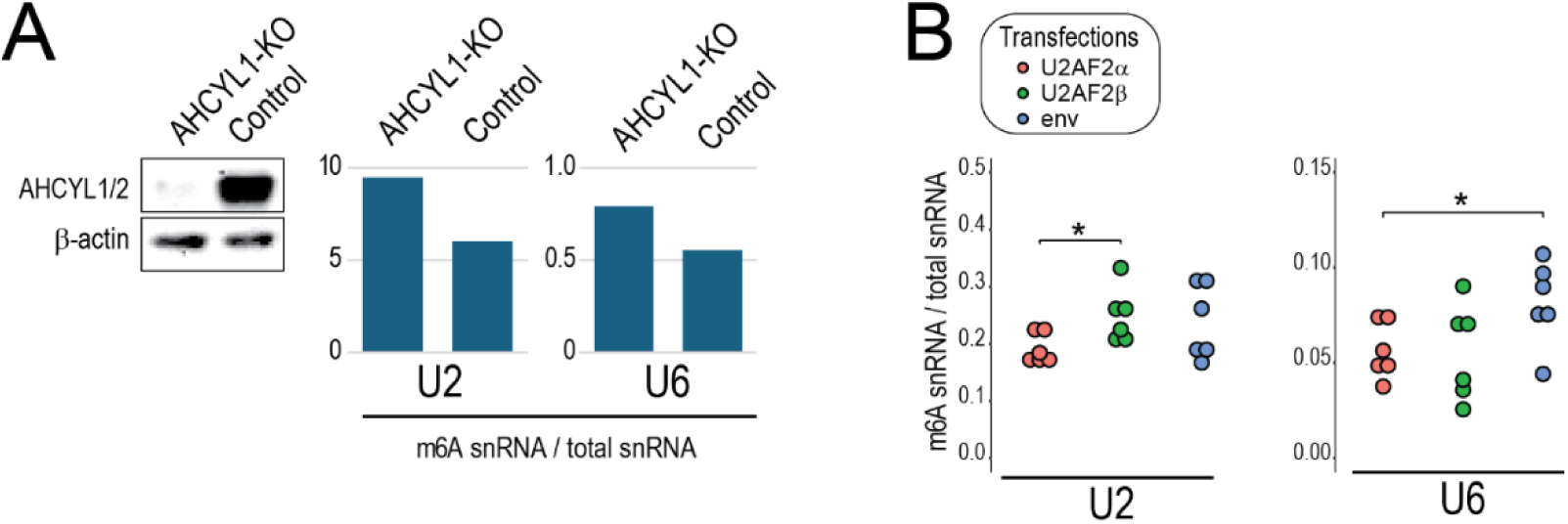
U2AF2 and AHCYL1 reduce m6A levels in snRNA. **A)** Impact of AHCYL1 knock-out. Western blot shows loss of AHCYL1 expression after inactivating the *ahcyl1* gene in HEK293T cells. Graphs show m6A levels measured in U2 or U6 snRNA, compared to the total snRNA amount. **B)** Impact of U2AF2 paralogues. The levels of m6A in snRNA were measured as in **A)**, in total RNA from cells transfected with *F. borealis* U2AF2 or env.

### Larvacean snRNA shows specific N^6^-methyladenosine modification patterns

As *F. borealis* U2AF2 paralogue could reduce m6A levels on mammalian snRNA, we considered the possibility that the snRNA of *F. borealis* could carry base modifications that differ from other species. We examined m6A levels in the snRNA from a variety of organisms, including the larvacean *Oikopleura dioica* and the ascidian *Ciona intestinalis*. The full set of major snRNA could be detected in all samples (**Fig 9A**), but signals corresponding to m6A modification were distributed differently in larvacean snRNA (**Fig 9B**). First, m6A levels on U6 are considerably decreased in *F. borealis* compared to other species. In yeast, nematodes and humans, the methyltransferase METTL16 is responsible for modifying A_43_ in U6 snRNA, and the loss of m6A in *F. borealis* U6 snRNA does not seem to be caused by the loss of methyltransferase activity, since the genome conserved all m6A writers with demonstrated roles in RNA modification^31^ (**Fig 9C**).

**Figure 9:**
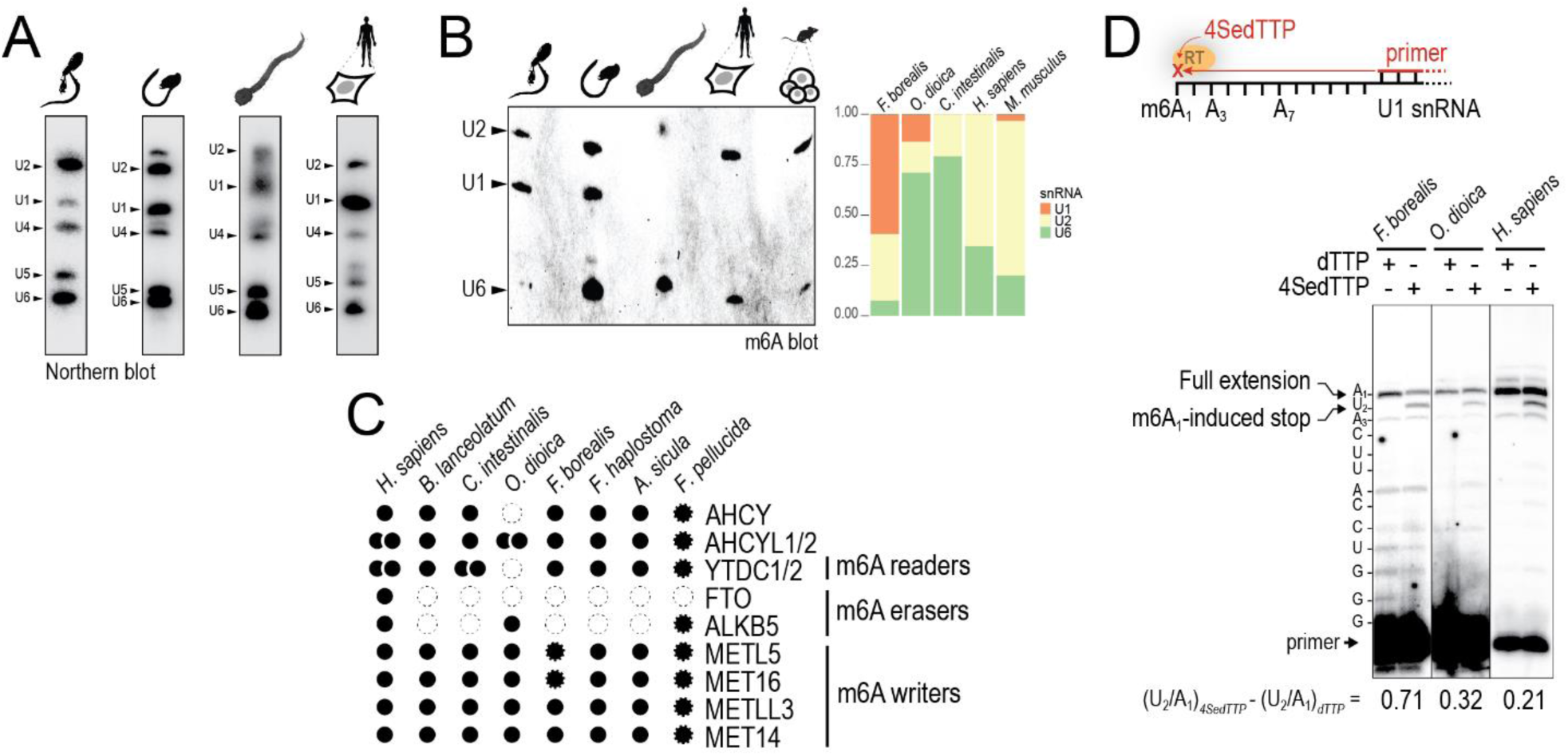
Divergent m6A patterns in *F. borealis* snRNA. **A)** Conservation of U2-type snRNAs. Northern blots show snRNA signals in total RNA extracted from (panels from left to right): maturing adults of *F. borealis* and *O. dioica*, *C. intestinalis* larvae and HEK293T cells. **B)** Detection of m6A. The immunoblot shows m6A signals in total RNA from the same origin as in **A)**. The last lane corresponds to total RNA from mouse ES cells. Bands were identified by comparing the migration profile with Northern blot experiments. The graph represents the proportion of signal intensity for U1, U2 and U6 snRNA. **C)** Conservation of m6A regulatory genes. We examined for each organism the number of copies of different genes involved in the SAM metabolism and its regulation (AHCY, AHCYL1/2), or in the deposition of m6A. Copies are represented by dots, full border indicates evidence gained with transcripts and genomic DNA, dashed border indicates evidence from genomic DNA only. Empty dots represent gene losses. **D)** Mapping m6A modification in the 5’end of U1 snRNA. Top, representation of the experimental approach showing that the presence of m6A on the RNA template induces reverse transcription (RT) stop when the thymidine analogue 4SedTTP is incorporated. Bottom, result of a primer extension experiment conducted on total RNA from three distinct species. We measured for each condition the ratio of cDNA stopped at position U_2_ of U1 snRNA relative to full-length cDNA. Values at the bottom of lanes correspond to the ratio measured in the presence of 4SedTTP minus the ratio measured with dTTP.

Therefore, we cannot exclude that m6A loss on U6 snRNA is caused by the gain of lineage-specific molecules, which could include U2AF2β. Second, both larvaceans showed prominent m6A signals corresponding to U1 snRNA, while using the same technique, m6A could be detected on mammalian U1 snRNA only in an FTO^-/-^ background^30^. Using a procedure recently made available for direct m6A mapping^32^, we confirmed that the signal detected with m6A immunoblot, corresponds to a m6A modification that is present at the first adenine of U1 snRNA in *F. borealis* and *O. dioica* (**Fig 9D**). Interestingly, we could not find an FTO homologue outside mammals, showing that in organisms like *C. intestinalis* where m6A modification on U1 snRNA is undetectable, other erasers could handle the control of m6A levels. Intriguingly, the expression of U1 snRNA appears lower in *F. borealis* compared to other species (**Fig 9A**). We cannot exclude that the lower signal could be related to the probe’s characteristics (labelling efficiency for instance), but it could also show that U1 snRNA plays a more limited role during pre-mRNA processing.

### Impact of m6A re-patterning for the recognition of 5’ss

The transcription start adenine A_1_ of U1 snRNA also bears a 2’-O-methyl in mammals^30^, and pseudo uridylations at present positions Ψ_4_ and Ψ_5_ in U1 snRNA can regulate splicing as they are directly engaged in the duplex with the 5’ss^33^. In the absence of a strong complementarity with the non-canonical 5’ss of *F. borealis*, it remains difficult to predict whether the 5’end of U1 snRNA will anneal to the same position on the pre-mRNA or if it would instead use a shifted register, as shown by others^34^. Because it has a stabilizing effect on base stacking^35^, unpaired m6A_1_ could help to form duplexes between U1 snRNA and non-canonical 5’ss (**Fig 10A**). The m6A present at this position could also regulate the transfer of the 5’ss from U1 snRNP to the U4/U6.U5 tri-snRNP when the pre-catalytic spliceosome is assembled. During this step, Prp28 unwinds the 5’ end of U1 snRNA, making the 5’ss accessible to interact with the ACAGAGA box of U6 snRNA^36^.

**Figure 10:**
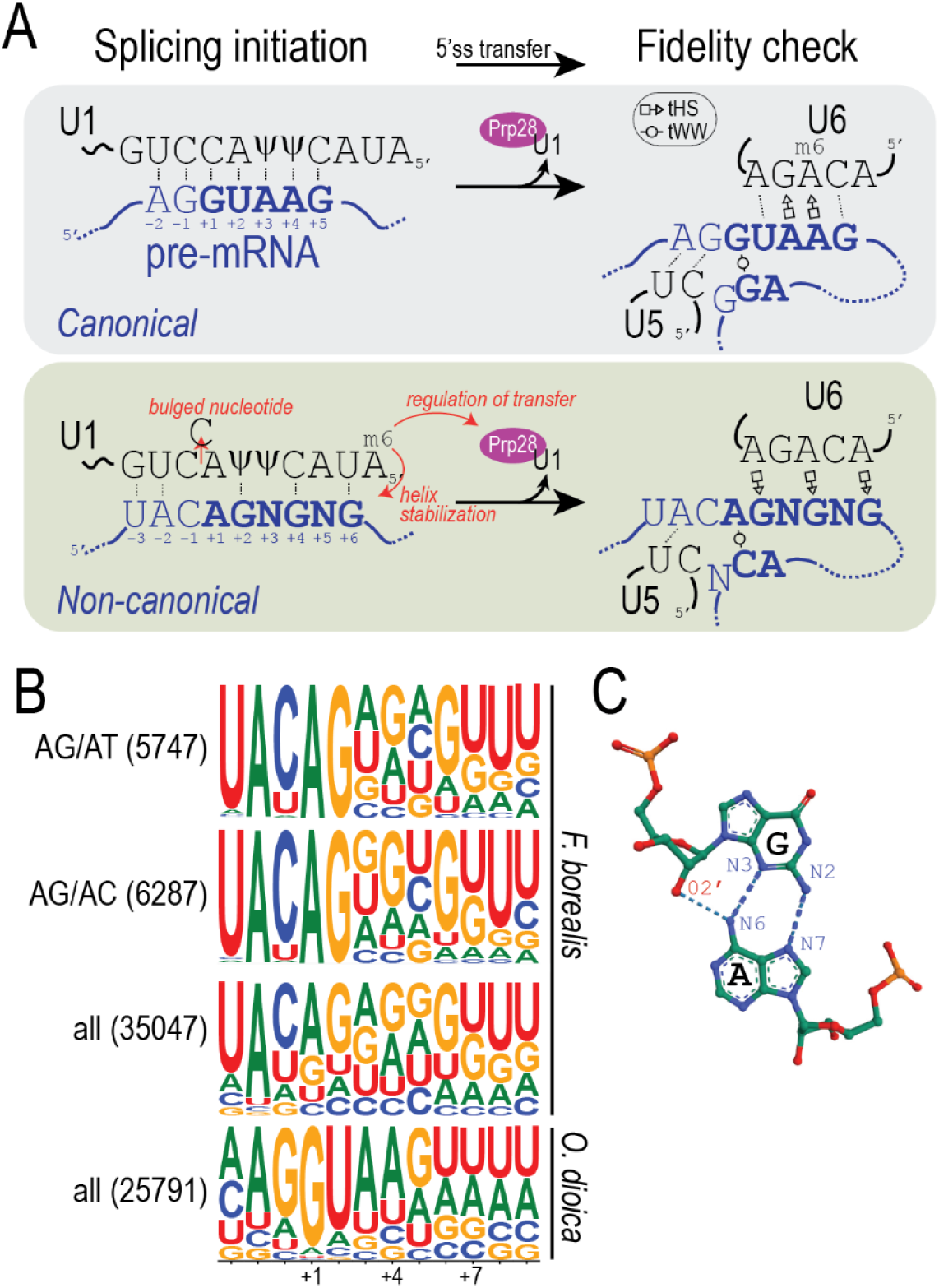
Function of m6A modification during non-canonical splicing. **A)** The RNA network of 5’ss recognition. The top panel is a model based on Artemyeva-Isman & Porter^7^ representing base-pair interactions established between the 5’ss (intronic residues shown in bold) and U1 snRNA during the initiation of splicing, when the U1 snRNP scans for exon/intron boundaries; and between the 5’ss and U6 snRNA after the pre-mRNA is transferred to the tri.snRNP. The assembly of an RNA network involving U5 and U6 snRNA on one hand, and intron termini on the other hand, constitutes a fidelity check that ensures splicing accuracy. Dashed lines represent Watson-Crick interactions, symbols represent non-Watson-Crick pairs. The bottom panel proposes a model of interactions established on a non-canonical 5’ss representing the major classes of *F. borealis* introns. Here, the binding of U1 snRNA may involve shifted register and stabilization by m6A_1_. **B)** Sequence logos of 5’ss showing overrepresentation of guanidine in *F. borealis* introns. **C)** Example of tHS interaction between the sugar edge of G_2657_ and the Hoogsteen edge of A_2664_ in the Sarcin/Ricin Domain from E. coli 23 S rRNA (PDB 3DVZ)^63^.

Compared to other species, adenines are less frequent beyond the position +2 of *F. borealis* introns and we see instead an over-representation of guanines (**Fig 10B**). This region is normally targeted by the m6A-modified ACAGAGA box, raising the possibility that the loss of conserved m6A in U6 snRNA is related to 5’ss divergence in this organism.

When interacting with the eukaryotic 5’ss consensus, the modified m6A_43_ in U6 snRNA establishes a non-Watson-Crick base pair with the adenine at position +4 of the intron^37^ under the *trans* Hoogsteen / Sugar geometry (tHS)^38,39^ (**Fig 10A**). Reducing m6A levels in the ACAGAGA box favors the splicing of 5’ss where a uridine is present at position +4, suggesting that another geometry can be accommodated in the duplex^37^. By aligning the U6 snRNA on the 5’ss consensus of the two main classes of *F. borealis* introns, we predict that tHS interactions can be established between adenines of the ACAGAGA box and the conserved guanines at positions +2, +4 and +6 of the 5’ss. The loss of m6A modification in the ACAGAGA box could play a critical role in stabilizing tHS pairing with guanine, by enabling an additional hydrogen bond between the N6 amine of A_43_ of U6 and the sugar of the opposite guanine on the 5’ss (**Fig 10C**).

## Discussion

By placing a variety of organisms under molecular scrutiny, genome projects conducted during the last decade have led to fundamental discoveries about the origins of introns and the evolution of the spliceosome. Important changes to the repertoire of spliceosomal genes were revealed in unicellular organisms, and these results have proven valuable for appreciating how splicing function can be preserved and how its mechanisms have adapted to intron variety^16,40^. Such knowledge is instrumental for deciphering splicing dynamics in the complex genetic environment of the human genome. By revealing how the spliceosome can change its selectivity, we prove that organisms genetically distant from mammals are still relevant for illuminating the function of conserved cellular machinery.

On a broader perspective, the results presented in this study open new perspectives about the evolvability of splicing across eukaryotes.

### The divergence of U2AF and its impact on 3’ss recognition

In this study, we showed that the divergence between the paralogues of *F. borealis* U2AF subunits affected their interactions with RNA and their ability to interact with protein partners. These changes have an impact on splicing function, as we could document that the presence of diverse *F. borealis* U2AF induce specific splicing events in mammalian cells. The function and RNA-binding properties of U2AF1 and U2AF2 have been extensively documented in mammals, and our results bring novel knowledge by showing how evolution can change the biochemistry of these crucial splicing factors. Documenting with biophysical approaches how the different U2AF complexes present in *F. borealis* interact with RNA, could reveal if the recognition mechanisms canonically employed by U2AF^41^ have been adapted to the intron landscape of *F. borealis*, and which mutations in U2AF1β and U2AF2β determine the divergent affinity for RNA. In both U2AF2 paralogues, the N- terminal region upstream the ULM corresponds to a RS domain rich in polar residues. The inhibitory effect on the RNA binding function in U2AF2β might be controlled by phosphorylation, as was shown for the C-terminal RS domain of SRSF1^42^. Although full-length U2AF2α showed a strong affinity for RNA, we cannot exclude that the RS domain also plays a regulatory function in this paralogue. Compared to U2AF2β, one remarkable difference in U2AF2α is a proline stretch between residues 133 and 137, which could reduce the flexibility of the disordered N-terminus and constrain its ability to interact with RRMs (**FigS5**). Further experiments employing mutant U2AF2 should figure out the impact of N-terminal domain phosphorylation and proline conformation on the affinity for RNA.

### Approaching the new functions of U2AF

Expression of U2AF genes *in situ* shows distinctive patterns in *F. borealis* (**FigS1**). The divergent paralogues are expressed in the germline, while the conserved paralogues are preferentially expressed in tissues like the tail muscle and gut cells. These patterns may reflect a sub functionalization, corresponding for instance to the control of tissue-specific splicing events. Deploying spatial transcriptomics in *F. borealis* will be instrumental for determining if the processing of different classes of introns is linked to cell type, and if it correlates with the expression of different versions of U2AF. Dissecting spliceosome functions in *F. borealis* is still a challenging task and in the short term, we believe that heterologous systems such as mammalian cell cultures will remain instrumental for gaining answers about non-canonical intron recognition. With genetic engineering to replace the endogenous U2AF genes with *F. borealis* U2AF1β and U2AF2β, we could gain a background-free system that is adequate to reveal the links between RNA binding and splice site selection.

By showing that U2AF2β binds AHCYL1 in mammalian cells, we revealed an unexpected link between splicing factors and regulators of the SAM metabolism. A variety of roles have been attributed to AHCYL1, but how it interferes with AHCY function remains largely unknown^26^. The protein sequence of AHCYL1 is remarkably well-conserved in tunicate genomes, which are otherwise characterized by frequent gene loss and remarkably high evolution rates^43^. It suggests a strong selective pressure for preserving AHCYL1 functions, and future studies should test its impact on animal development and gene expression.

### Evolutionary and chemical constraints to the selectivity of the spliceosome

Most components of the human spliceosome are conserved in intron-rich yeasts, but cases of gene loss were reported in several lineages of unicellular eukaryotes^16,17^. For these organisms, the loss of snRNP genes appears to be linked to reduction of intron diversity, often in a context of rapid evolution. These studies have been instrumental for revealing the function of genes that have been lost during evolution, and they recently showed that the massive reduction of splice site diversity observed in Saccharomycotina genomes coincided with the loss of non-essential proteins that enforce splicing fidelity by repressing the usage of cryptic splice sites^16^. Nematodes and larvacean tunicates are the only metazoan phyla that have lost all snRNA components of the U12-type spliceosome, and for these genomes where a substantial number of non-GTAG introns remain, there is no apparent link between the loss of spliceosome genes and the diversity of splice sites^4,10,43^. We suppose that losing the U12-type equipment can impact the ability to process a larger variety of introns, which in the long term could compromise the genome’s resistance to splice site erosion or to invasions of introner transposons carrying weak splice sites^10,11^. A viable solution during evolution would be to compensate the loss of flexibility in the splicing machinery by introducing new components, or by modifying the function of existing ones.

Mutation of essential genes such as U2AF1 and U2AF2 can compromise cellular functions, and the gain of extra copies also bears the risk of giving rise to negative dominants. By analogy to essential splicing factors that have a functional homologue in the minor spliceosome, the risks associated with maintaining U2AF duplicates could be outweighed by a critical need for removing non-canonical introns. Retrospectively, the case of *F. borealis* U2AF appears as a common example of neofunctionalization mediated by gene duplication. In this organism, the situation of U1 snRNA genes appears more subtle. Here, the answer to the massive change of 5’ss identity was not evolving U1 snRNA sequences adapted to new introns - *F. borealis* has two U1 snRNA genes whose 5’ ends are strictly identical^10^; but instead, changing the basal level of m6A_1_ in U1 snRNA. This could be an advantage, as the possibility to duplicate and evolve divergent copies of U1 snRNA could be constrained to a much higher level by the critical role of U1 during splicing. The presence of m6A in mammalian U1 snRNA has been examined in a handful of studies only, and the results gained in our study open the exciting possibility that the regulation of this modification could have an impact on U1 function and splice site selection. The effect of m6A_1_ on U1 snRNA function during splicing should be examined with adequate experimental design because on its own, transcriptome analysis of FTO^-/-^ cells cannot permit to isolate the effects caused by the presence of m6A_1_ in U1 snRNA^30^, from those caused by the loss of m6A on pre-mRNA sequences^44^.

We propose that the large diversity of intron ends present in the *F. borealis* genome ultimately reflects the inherent flexibility of the spliceosome’s catalytic site, which can potentiate the effect of snRNA base modifications during splice site selection. Splicing depends on the conservation of non-Watson-Crick base pairing between the terminal nucleotides of introns^7^. The isostericity of G-G and A-C terminal pairs (respectively formed with GT/AG and AT/AC introns) has been raised as the molecular basis to explain why the major spliceosome can process AT/AC introns in laboratory experiments^6^. Based on mutant analysis^45^, G-G and A-C intron termini were predicted to adopt a *trans* Watson-Crick / Watson-Crick (tWW) conformation in the human spliceosome. In *F. borealis*, despite the extreme variety of splice sites, the termini of at least 55% of introns could form tWW interactions isosteric or nearly isosteric to G-G *i.e.*, with isodiscrepancy indexes under 3.5^46^ (**TableS2**). If we consider an alternative *cis* Watson-Crick / Hoogsteen (cWH) conformation that is supported by CryoEM data of the *S. cerevisiae* spliceosome^8^, this proportion will correspond to 45% of *F. borealis* introns. Examining the presence of other base modifications on the snRNA and on the pre-mRNA and ultimately, high-resolution experimental structures of the *F. borealis* splicing machinery will give definitive answers to this hypothesis.

## Acknowledgments

Simon Henriet acknowledges financial support from the Norwegian Cancer Society (Pioneer grant #227930 – “Non-canonical splicing as a source of aberrant genetic information and genomic”). We thank Ave Tooming-Klunderud at the Norwegian Sequencing Centre in Oslo for help during Pacbio sequencing experiments. We thank Teshome Tilahun Bizuayehu and Jonas Brandenburg at the Michael Sars Centre for help during ONT sequencing experiments. We thank the Norwegian Research and Education Cloud for generous access to computing infrastructure. Larvacean samples were provided by Anne Aasjord, animal facility manager at the Michael Sars Centre. Ascidian samples were provided by the Chatzigeorgiou lab at the Michael Sars Centre. Mouse ES cells were provided by Max Jordi Makem Pekouankouang at the Michael Sars Centre.

## Material and Methods

### Animal samples

Specimens of *Fritillaria borealis* were sampled from laboratory populations established with specimen collected near Bergen, on the west coast of Norway^15^. Specimens of *Fritillaria haplostoma* and *Appendicularia sicula* were collected directly near Bergen, on the west coast of Norway^10,15^. Specimens of *Fritillaria pellucida* were collected near La Jolla, (CA, USA)^10^.

### Genome sequencing

We extracted high-molecular weight genomic DNA from mature *F. borealis* gonads dissected under microscope, with the Chomzynsky procedure. The DNA corresponding to 190 individuals was pooled, and 1.5 µg was used for long-read sequencing with Oxford Nanopore Technologies (ONT) platforms. We prepared ONT libraries following the manufacturer’s recommendations (Ligation sequencing DNA V14, ONT) but increasing incubation times for adapter ligation to one hour and bead elution time to 14 hours to improve DNA recovery. Between 250 and 300 ng of ONT library were sequenced on a MinION flow cell (R10.4.1). Available pores rapidly decreased during the run, presumably due to pore blocking by polysaccharide contaminants present in DNA preparations. To get enough coverage for genome assembly, we collected over 29.10^4^ reads with N50=4.9 kb produced by sequencing three independent libraries.

The DNA from whole bodies of four mature *F. haplostoma* individuals was prepared as described above and prepared for Illumina sequencing (Nextera XT DNA Library Preparation Kit). These libraries were barcoded and pooled with *A. sicula* libraries prepared for an earlier study, using whole-genome amplified DNA (WGA-DNA) obtained from individual specimens^10^. Pooled libraries were sequenced with 150bp PE on a 1.5B NovaSeq X flow cell at the Norwegian Sequencing Centre (NSC, Oslo, Norway), producing 56.10^7^ and 87. 10^6^ reads for the *F. haplostoma* and for the *A. sicula* libraries, respectively. We prepared ONT libraries for *A. sicula* using WGA-DNA produced from single individuals, as described above. We sequenced 400 ng of *A. sicula* ONT library on a PromethION 2 solo flow cell (R10.4.1) for 16.5 hours, producing 2.10^6^ reads with reads N50=6.4 kb.

### Transcriptome sequencing

We extracted total RNA (Nucleospin RNA XS, Macherey-Nagel) from *F. borealis* (24 larvae, 3 immature and 8 mature adults), *F. haplostoma* (6 mature adults) and *A. sicula* specimens (one mature adult). Individual transcriptomes were converted to cDNA and amplified with barcodes for a total of 17 cycles, following Pacbio’s guidelines for isoform sequencing (Iso-Seq Express Template preparation). SMRTbell libraries were prepared from pools of barcoded cDNA and sequenced with two SMRT cells on a Sequel II instrument at the NSC. Circular Consensus Sequencing (CCS) reads and Full-Length Non-Concatemer reads were produced with the Iso-Seq pipeline (SMRT Link v10.2).

### Genome assembly

We used long-read sequencing data to improve an existing assembly of the *F. borealis* genome produced with short-read data from a single individual^18^. We processed ONT reads with Guppy software v6.4.8 configured for GPU-accelerated basecalling with superior accuracy model (dna_r10.4.1_e8.2_260bps_sup). We assembled ONT reads using Flye v2.9.4, and we merged long- and short-read assemblies together using Quickmerge v0.3.

The hybrid assembly was polished with Illumina reads using Pilon v1.24, resulting in a last version with 4063 contigs, for a total size of 79.7 Mb and N50=39.9 kb.

To produce an *A. sicula* genome assembly, we used sequencing information obtained from WGA-DNA corresponding to a single individual. ONT reads were basecalled with Dorado V0.8.3, configured with superior accuracy model. We assembled ONT reads with Flye v2.9.4 and polished the assembly with Illumina reads produced from the same individual. The final assembly contains 13.10^3^ contigs, for a total size of 65 Mb and N50=6.9 kb.

We assembled 75.10^6^ paired-end Illumina reads from the *F. haplostoma* samples with Megahit assembler^47^. Heterozygous contigs were removed from the assembly with the Redundans pipeline, using the initial set of Illumina reads for scaffolding and gap-closing steps. The final assembly contains 55.10^3^ contigs, with a total size of 120 Mb and N50=2.2 kb.

### Gene and intron annotations

Gene evidence and intron positions were obtained by aligning Pacbio transcripts to the genome assemblies. Only high-quality Pacbio isoforms were employed, corresponding to clusters produced by Full-Length Non Concatemer (FLNC) reads that belong to the same barcode.

For *F. borealis*, we first used Cutadapt v1.16 to remove the splice leader sequence present at the beginning of transcripts^10^. We mapped a collection of 249123 trimmed transcripts on the genome assembly with minimap2, without an alignment guiding option that uses conservative positions flanking canonical splice sites (--splice-flank=no). We produced 34034 unique isoforms after collapsing transcripts with Python scripts^48^. After aligning transcripts to the genome, gap intervals were used to determine intron positions. We first collected 28845 introns whose borders could be aligned without ambiguity, corresponding to situations where the last position of the upstream exon and the first position of the downstream exon have different bases. In these non-ambigous introns, we could detect 88% of the time an adenine at the penultimate position, consistent with the strong conservation observed in various intron classes of other eukaryotes^4,7^. To precisely map 6203 additional introns with ambiguous borders, we applied a conservative set of rules : #1, intron ends must have terminal dinucleotides GT/AG; #2, introns that do not comply to #1 must have terminal dinucleotides AG/AC or AG/AT; #3, introns that do not comply to #2 must have a penultimate adenine.

To annotate introns in the *F. haplostoma* and the *A. sicula* genomes, we used PacBio’s Iso- Seq package to align transcripts over genome assemblies and to collapse redundant isoforms. As described above, we first established intron borders based on splice signals discovered with a set of non-ambiguous introns positions (1507 and 1381 introns for *F. haplostoma* and *A. sicula*, respectively). For both genomes, the following rules were applied to map additional introns with ambiguous borders: #1, intron ends must have terminal dinucleotides GT/AG; #2, introns that do not comply to #1 must have terminal dinucleotides GC/AG; #3, introns that do not comply to #2 must begin with either GC/AG/AT/GG and end with either AG/AC/AT; #4, introns that do not comply to #3 must have a penultimate adenine.

To annotate introns in the *F. pellucida* genome, we first used BLASTX to find matches against a collection of 30987 proteins predicted from the *F. borealis* isoform collection. Intron borders were found based on gap intervals. We kept 450 non-ambigous cases for which no gaps are present and amino-acid identity is strictly conserved in at least two codons out of five, in each flanking exon.

### Homology search

Gene orthologues were annotated with reciprocal blast. First, we used human protein sequences as queries against transcriptomes and genome assemblies of the species of interest. We searched for orthologues in the annotated transcriptome and in the genome assembly of *O. dioica*^43^ and for other larvaceans, we employed the genomic resources described in this study. Orthologues from other organisms were searched in the ENSEMBL and ANISEED^49^ databases. Top hits were checked for protein length and domain conservation, then used as queries for reciprocal BLAST against the UNIPROT database to confirm annotation.

### Expression and purification of U2AF heterodimers

The full-length open-reading frames (ORFs) of U2AF subunits were amplified from the cDNA of *F. borealis* with Taq polymerase, and A-tailed PCR products were cloned in pGEM- T vector for sequencing. We employed a seamless cloning strategy (In-Fusion HD, Takara) to subclone U2AF ORFs in bacterial expression vector. We first replaced positions 100-168 of MCS-1 in pRSFDUET-1 with the U2AF2 ORF in frame with the 6xHis tag. Then, we replaced the whole MCS-2 with the U2AF1 ORF, in frame with an N-terminal tag that consists of 30 amino acids of the Twin-Strep-tag, followed by amino acids 2-56 of the B1 domain of Streptococcal Protein G^50^. Deletion mutants were engineered with seamless cloning.

We transformed tandem expression construct in *E. coli* BL21-AI. Clones were grown to OD=0.8 in terrific broth (TB) and protein expression was induced with 50 µM IPTG, in the presence of 0.4% L-arabinose and 0.1 M ZnCl_2_ during 16 h at 15°C. Cell pellets were lysed on ice in HHS buffer (HHS: Hepes 20 mM, NaCl 0.5 M, 10% glycerol, 1% Tween-20, pH 7.8) supplemented with 2 mg/mL lysozyme, 1 mM TCEP, 1 mM PMSF and 1x Protease Inhibitor Cocktail (cOmplete, EDTA-free; Roche). Lysates were sonicated on ice three times with 10 second pulses, then incubated 30 minutes on ice with DNase I, 2U/mL RNase cocktail (Ambion), and 3% BioLock (IBA). Lysates were centrifuged at 18000 g for 30 minutes at 4°C, and protein complexes assembled with U2AF1 were captured on a Strep-Tactin XT 4 flow gravity flow columns (IBA). Column-bound complexes were washed with 20 column volume (CV) of HHS, 5 CV of HLS buffer (HLS: Hepes 20 mM, NaCl 0.1 M, 5% glycerol, 0.5% Tween-20, pH 7.8), and 10 CV of HLS buffer supplemented with 5 mM biotin. Purified complexes were eluted 6 times with 1 CV of HLS supplemented with 0.1 M biotin. We performed buffer exchange on pooled fractions by centrifugation on 10 kDa MWCO columns (Zeba Spin, Thermo) equilibrated with HLS buffer.

### Electrophoretic mobility shift assays

We used seamless cloning to insert the EMSA probe sequence in pUC19 downstream a T7 promoter and immediately upstream a SapI restriction site. Constructs were linearized with Sap I and used as templates for in vitro transcription (IVT) reactions (MEGAshortscript T7 transcription kit, Ambion). IVT products were treated with Turbo DNase (Ambion), purified with Phenol/Chloroform extraction and treated with Calf Intestinal Alkaline Phosphatase (Thermo). Dephosphorylated probes were purified on RNA Clean & Concentrator-5 (RNA-CC5, Zymo Research). We end-labelled 100 ng of probe with T4 PNK (NEB) and 10 µCi [γ^32^P]-ATP at 6000 Ci/mmol, and reaction products were purified on 8% PAGE/TBE-Urea 8M. For EMSA, 100 cpm probe were denatured 2 minutes at 80°C, placed on ice, and incubated 1 h on ice in EMSA buffer (HLS supplemented with 50 ng/µL BSA, 1% RNase inhibitor (Protector, Roche), 0.5 mM TCEP, 0.5 mM EDTA) in the presence of purified U2AF. RNPs were run on 6% PAGE/TBE 0.5x for 2 h at 4°C under 5V/cm. Gels were fixed with a 10% EtOH/5% AcOH solution and dried under vacuum. We recorded signals with phosphor screens and a Typhoon FLA scanner (GE), and measured signal intensity with ImageJ. Dissociation constants were determined with the nlsfit function of the R package easynls, using an equation for one-site model: Kd= [P][R]/[PR] where P and R are the free U2AF and probe, respectively, and PR is the U2AF-probe complex. Our conditions provide at least 5-fold molar excess of U2AF compared to the probe.

### RNA-compete assays

We designed a pool of 5633 unique sequences derived from *F. borealis* introns, which include the 20 last nucleotides of the introns, followed by 10 nucleotides of the downstream exon. The T7 promoter sequence followed by an extra guanine residue, was added at the 5’ end of the pool. A SapI restriction site was added at the 3’ end, followed by a target sequence for PCR priming. After chemical synthesis (Twist Biosciences), we produced IVT templates from oligo pools with PCR reactions and primers specific to intron-flanking sequences. To preserve complexity and to avoid the formation of unspecific products, amplification rounds were performed with at least 60% of the initial pool as template during a maximum of 6 PCR cycles. Template linearization, IVT reactions and 5’ phosphate removal was performed as described above. We checked the integrity of RNA pool on 15%PAGE/TBE-Urea 8M.

We used seamless cloning to insert the full-length ORF of U2AF1 in pGEX-6P-1, in frame with the N-terminal GST. Constructs were transformed into *E. coli* BL21, selected clones were grown to OD=0.8 in TB and protein expression was induced 16 h at 18°C with 0.4 mM IPTG. Cell lysis and nucleic acid removal was performed as described above, in LY buffer (LYB: 25 mM Hepes, 0.35 M NaCl, 10% glycerol, 0.5% Tween-20, pH 7.5). After 10000 g centrifugation for 30 minutes at 4°C, fusion proteins were captured on Glutathione-sepharose 4B (Cytiva) for 2 h at room temperature, washed with 30 bead volume of LYB and eluted 4 times with 1.5 bead volume of LYB supplemented with 50 mM glutathione. We exchanged the elution buffer to LYB as described above.

For RNA-binding assays, 2 picomoles GST-U2AF1 were bound to 2 µL Glutathione High-Capacity Magnetic Agarose Beads (Millipore) for 1 h at room temperature in 0.2 mL of RB buffer (RBB: 25 mM Hepes, 0.5 M NaCl, 5% glycerol, 1 mM EDTA, 1 mM DTT, 2 mM MgCl_2_, 0.5% Tween-20). After three washes with RBB to remove unbound proteins, we added 1 µM of RNA pool to the beads in a final volume of 0.5 mL RBB, together with 10 µg BSA, 4 µg Heparin, and 2 µL of RNase inhibitor. Samples were incubated 2 h at 4°C on wheel, and beads were washed three times with 0.5 mL ice-cold RBB. Proteins were digested with Proteinase K in 10 mM Tris, 1% SDS, 2 mM EDTA for 15 minutes at 37°C, and we recovered RNA with RNA-CC5. We prepared Illumina libraries using NEBNext Small RNA library prep kit (NEB), and samples were sequenced on a MiSeq instrument at the NSC. After adapter trimming, an average of 84% reads could be mapped on the original collection of introns with Bowtie2 (--very-sensitive-local).

We analyzed an average of 945.10^3^ reads per library. We filtered out introns with low read count (no sample with >10 reads). Next, read counts were normalized against the total number of reads per sample and for each intron, we measured enrichment using: 𝐸 =log[(C1+2)/(C2+2)], where C1 and C2 correspond to read counts measured for different U2AF1 paralogues. We first selected introns with E>0.12 when comparing differential enrichment between either *F. borealis* U2AF1α and U2AF1β, or *O. dioica* U2AF1 and F. borealis U2AF1β. We scored introns with their read count, and we selected a group of top-scoring introns representing 80% of the total read count kept for sequence analysis. Composition statistics and motif detection were performed on RNA pool sequences clipped from their terminal dinucleotides.

### Mammalian cell expression and transfection

We used seamless cloning to introduce *F. borealis* U2AF2 paralogues in pCDNA3.1, in N- terminal fusion with a 6xHis tag. U2AF1 paralogues were introduced in pSF-CMV, in fusion either with a N-terminal GST, or with the Twin-Strep tag and solubilization domain as described above. Constructs were delivered to HEK293T cell grown in DMEM using Lipofectamine 3000 (Invitrogen), and cultures were harvested 48 h post-transfection.

### Long-read sequencing of cDNA

Cells transfected with U2AF1 and U2AF2 constructs were treated with 0.15 M cycloheximide for 8 h prior harvest. Controls were treated with 0.1% DMSO. Cells pellets were washed in PBS, and RNA was extracted with TRIzol reagent (Invitrogen), following the manufacturer’s procedure. Samples were treated with Turbo DNase, and 1 µg RNA was converted to cDNA with Superscript IV reverse transcriptase, using a combination of random 10-mers and oligo-dT primers. For each target gene, we amplified cDNA with specific primers in separate PCR reactions, for a total of 20 cycles. PCR products from the same cDNA were pooled together and purified with AMPure XP SPRI beads (Beckman). We prepared libraries with barcodes to separate cDNA samples, following ONT’s procedure for barcoding and sequencing amplicons (SQK-LSK114). We sequenced 55 ng of library pool on a PromethION 2 solo flow cell (R10.4.1) for 2 h, producing 6.10^6^ reads with N50=3.4 kb. Reads over 500 bp with quality score over Q9 were basecalled with Dorado V0.8.3, configured with high accuracy model. We trimmed adapters and primers with porechop, and processed the reads with TALON v6.0^51^, with a database corresponding to regions of the human genome targeted by the gene-specific primers. For each sample, we considered only isoforms whose sequencing coverage across samples is superior to 50, and we found splicing events based on exons positions mapped by TALON. Introns were classed as canonical (ending with GT/AG) or non-canonical (other combinations of terminal dinucleotides). Isoforms were classed as canonical unless their splicing involved at least one non-canonical splice site.

For each isoform of each target gene, we scaled expression across samples with: E= (C - Mc)/Sc where E is the scaled isoform expression value in the sample, C is the isoform read count in the sample, Mc is the mean of the isoform read count across samples and Sc is the standard deviation of the isoform read count across samples. For each sample, we collected intron positions corresponding to the splicing of isoforms with E > 0.75, and we established a list of specific positions absent from other samples. We analyzed sequence composition using a non-redundant collection of introns. The PPT score of each intron was calculated with the rules established by Clark & Thanaraj^52^.

### Transcriptome analysis of U2AF2-expressing cells

We used Trizol to extract total RNA from HEK293T cells transfected with either U2AF2 constructs described above, or with the pCDNA3.1 vector containing the env gene of a LTR retrotransposon^19^. We enriched mRNA with the Dynabeads mRNA Purification kit (Ambion) and barcoded sequencing libraries were prepared from poly-A^+^ RNA with NEBNext Ultra II RNA Library Prep kit (NEB). Pooled libraries were sequenced 150bp PE on a 1.5B NovaSeqXPlus flow cell at the NSC.

Basic trimming of raw reads was performed with FastP v0.24.0 to remove adapter sequences and keep Q20 reads^53^. The data was mapped with STAR v2.5.2b^54^ to *H. sapiens* GRCh38.p14 primary genome assembly using the latest gene annotation file retrieved from GENCODE^55^. The mapping results were used to run an additional STAR two-pass mapping for further analysis of AS events. We used Salmon v1.10.3^56^ to quantify gene expression and we conducted Differential Gene Expression (DGE) analysis with R packages tximport v1.32.0 and DeSeq2 v1.44.0^57,58^. Differentially expressed genes (DEGs) were filtered with significance threshold of -0.58 ≤ Log2FoldChange value ≥ 0.58 and adjusted p value padj < 0.05. To examine enriched functional categories and pathways in DEGs, we performed GO and KEGG Over-representation analysis (ORA) using the clusterProfiler R package v4.12.6^59^. We determined AS events with two-passed STAR genome aligned bam files using the rMATS turbo software v4.3.0^60^ with --novelSS function enabled to allow the detection of novel splice sites. The AS events detected were filtered using a significance threshold of FDR ≤ 0.05.

### Identification of U2AF2 partners

Cells transfected with either U2AF constructs or a control containing the *env* gene were washed in PBS and lysed in HLS, supplemented with 1 mM TCEP, 1 mM PMSF, and protease inhibitors (cOmplete, Roche). Protein concentration was determined with bicinchoninic acid assay, and for each transfection, a sample equivalent to 6 µg protein was used for immunoprecipitation. Samples diluted in 1 mL HLS were first pre-cleared by incubation with Anti-HA magnetic beads (Sigma, equivalent to 70 µL suspension) for 1 h at 4°C on wheel. The supernatant was transferred to Anti-V5 magnetic beads (Sigma, equivalent to 70 µL suspension) and incubated 2 h at 4°C on wheel. Beads were washed five times with 1.5 mL ice-cold HLS and resuspended in Laemmli buffer. Bound proteins were eluted for 5 min at 70°C and separated on Bis-Tris SDS-8% PAGE. After Coomassie staining, bands between 37 and 100 kDa were excised. Peptides were digested in-gel and analyzed on an Orbitrap Eclipse Mass Spectrometer at the Proteomics Unit of the University of Bergen. For validation of protein interactions, samples separated on gel were transferred to a PVDF membrane and tested with either SAHH Antibody (sc-271389, Santa Cruz Biotechnology) or AHCYL1/SAHH-3 Antibody (sc-271581, Santa Cruz Biotechnology).

### Generation of AHCYL1 KO cells

We transfected HEK293T cells with a set of CRISPR/Cas9 KO plasmids directed against the human AHCYL1 gene (sc-417109, Santa Cruz Biotechnology). Three days post-transfection, cells were harvested and washed in PBS supplemented with 2% BSA and 1 mM EDTA. Fluorescent cells expressing GFP and the Cas9 gene were sorted on a Wolf G2 benchtop cell sorter. Gating was set by examining the fluorescence of cell cultures transfected with pEGFP-N1 or with a LacZ-expressing construct. After sorting, GFP-positive cells and GFP-negative were pelleted, resuspended in complete culture media and grown four days at 37°C before assaying AHCYL1 expression and m6A levels.

### Detection of N6-methyladenosine in snRNA

Total RNA was extracted with Trizol, following the manufacturers’ recommendations. To examine m6A levels, total RNA was separated on 10% PAGE-Urea and transferred on a positively charged nylon membrane (Hybond N^+^, Amersham). We detected N6- methyladenosine using a m6A antibody (ab151230, Abcam). To quantify total snRNA amounts, we performed Northern Blot using a pool of radioactive, end-labeled oligoribonucleotides probes complementary to U1, U2, U4, U5 and U6. Membranes were hybridized to probes in ULTRAhyb-Oligo (Ambion) during 14 h at 42°C, then washed twice at 42°C with 2x SSC / 0.5 % SDS. Signals were recorded and quantified as described above. When measuring cellular m6A levels, each RNA sample was tested with m6A antibody and ran in parallel for Northern Blot. The m6A signal was quantified with Image Lab (Bio-Rad) and normalized against the intensity of the corresponding radioactive signal.

### Mapping of m6A in U1 snRNA

We designed primer extension assays based on procedures described by Hong et al.^32^. Briefly, 1.5 µg total RNA was annealed to 150000 cpm of gel-purified DNA primer complementary to nucleotides 13-33 of U1 snRNA and 5’ end-labelled with [γ^32^P]-ATP. After 2 minutes at 90°C, hybridization mix were placed 2 minutes at 50°C, and 10 U Superscript IV reverse transcriptase (Invitrogen) were added in RT buffer (50 mM Tris-HCl, 4 mM MgCl_2_, 10 mM DTT, 50 mM KCl, 30 µM dATP, 30 µM dGTP, 30 µM dCTP, pH 8.3) in the presence of 30 µM of either dTTP or 4SedTTP (Jena Biosciences). After 20 minutes incubation at 50°C, cDNA synthesis was stopped by addition of Urea gel loading buffer, and denatured elongation products were resolved on 20% PAGE-Urea sequencing gels. Signal was recorded on fixed gels and processed as described above.

