## Supplemental Figures and Tables for "Deployment of non-canonical splicing in tunicate genomes is mediated by divergent U2AF function and re-patterning of snRNA m6A modification"

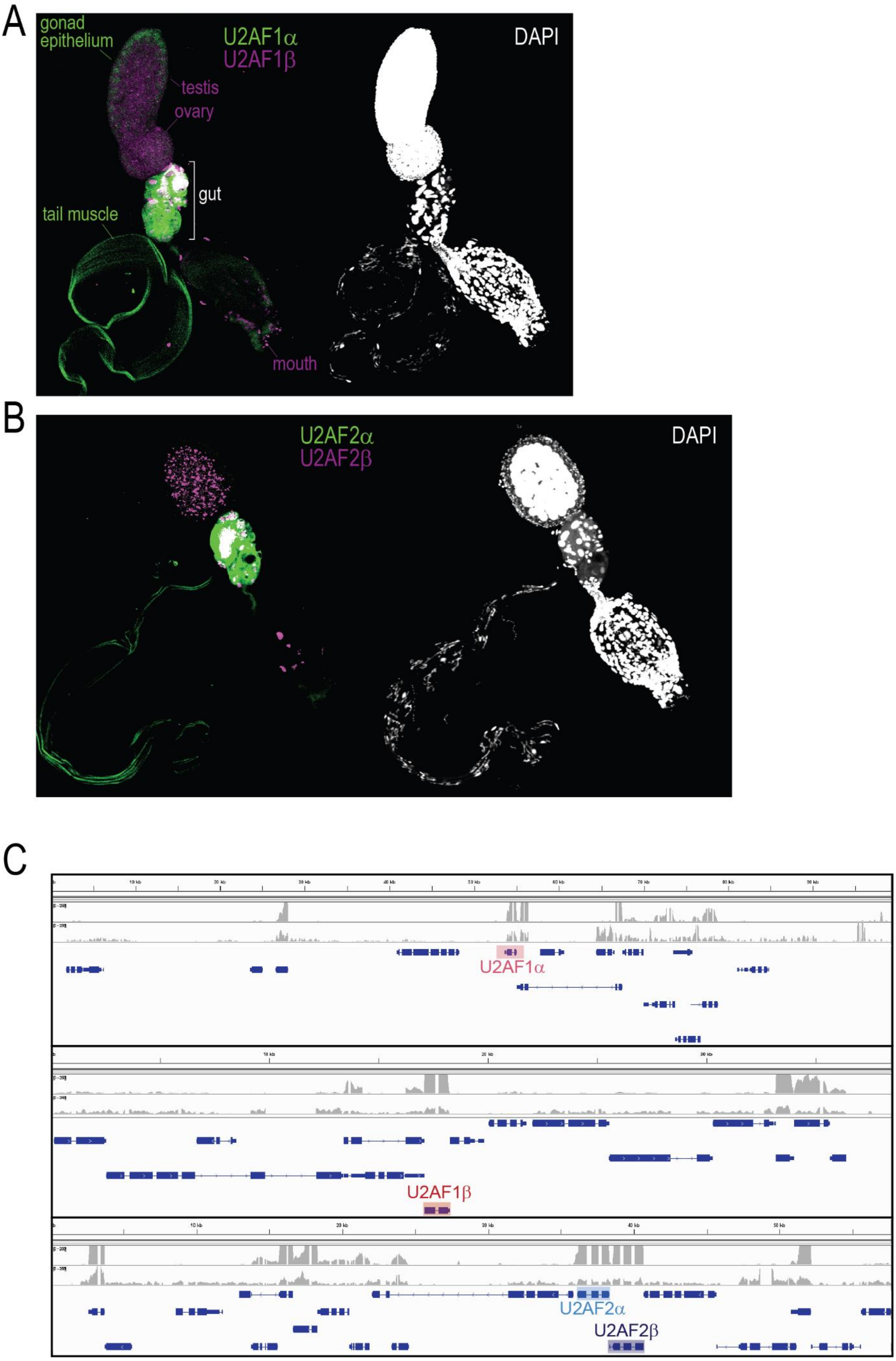
A
gonad epithelium
U2AF1α
U2AF1β
testis
ovary
gut
tail muscle
mouth
DAPI
B
U2AF2α
U2AF2β
DAPI
C
U2AF1α
U2AF1β
U2AF2α
U2AF2β

**<u>Figure S1:</u>** Expression of U2AF paralogues in *F. borealis*. **A)** Whole-mount HCR *in situ* hybridization performed on maturing adults, with fluorescent probes targeting either U2AF1$\alpha$ (green) or U2AF1$\beta$ (magenta). Nuclei were simultaneously stained with DAPI (white). **B)** Same as **A)**, with fluorescent probes targeting either U2AF2$\alpha$ (green) or U2AF2$\beta$ (magenta). **C)** Genome browser screenshots showing the gene models for U2AF1 and U2AF2 paralogues and their RNA-seq coverage during early or late developmental stages.

```
U2AF2A_Fbor        MSRNIKTETGEDADFEFDPDMDPSAYKLEPDVITTAPIKKQSRDYDRDSSNRHSSSRRES  60
U2AF2B_Fbor        MGKKDSDS-ESSGRDEYGIKVDPLDVKD---FGADAEVNRRKREKEDR------KKRRAR  50
U2AF2_Fpel         MPRNDSDS-ESSGRDEYGIKVDPADVKD---FGSHALYDEKKREKERE------RKRRER  50
                   * :: . .  ...  *:. .:**   *    . : *  ..:.*: :         .**

U2AF2A_Fbor        SSRRSESSRSSRSGRDRHDDRDSHRSRDSRSSRDSRDSRSHRDRSPRRDRDRRDRNRD--  118
U2AF2B_Fbor        SSSSSDDG-------HGHSARRKHRRRRRRSRSRSRDKSRKRRRSRDRSGDRNDRKRDRS  103
U2AF2_Fpel         RSRSSSSS--SSSSSSDDDRRRRKRRRRRRSRSRSRDRRRRKHRKRRRSS---------V  99
                    *  *...         .. *  :* *  **   ***   :: *.  *.

U2AF2A_Fbor        --GGSS--KVTFEEKE-------ASPPPPPRVRVNKYWDVAPEGFEHVQPNQYKEMQQNG  167
U2AF2B_Fbor        ASGSRSTSPIILKKKDSKKVRKRKEYWAWRNNNKFSFWDKAPEGFEHVTVKQYKEMQSTG  163
U2AF2_Fpel         DRSSRS---------------------KKPKVKKYKFWDIAPEGFEHVTVKQYKEMQATG  138
                     .. *                        . .  .:** ********  :****** .*

U2AF2A_Fbor        QLPIHFPGVVPGAAIAAQFPMAGGHVARQARRLYVGGIPFGATEQNMMEFFNAQMRTAGL  227
U2AF2B_Fbor        QVAVQSIGSHISGNID-EIKAGGSKNNRTYLSLYVGGLPFGATEQTVFEFFNAQMRAAKL  222
U2AF2_Fpel         QVAVQAPGATVSGNID-EVTRGGSKNNRTYLSLYVGGIPFGATEQNMGKVF---------  188
                   *: ::  *   .. *  :.  .*.:  *    *****:*******.: :.*

U2AF2A_Fbor        SQAPGDPILAVQINMDKNFAFLEFRSVDETTQALAFDGIQFMGQSLKIRRPSDYKAKPGQ  287
U2AF2B_Fbor        NDKTKAPIQAVRINMDKNFAFLEFKERSECTNALAFDGVEFRGAPLKIRRPLDYNPPSDD  282
U2AF2_Fpel         -------------NILXNFAFIEFKERAECTNALAFDGVEYRGAPLKIRRPLDYKPPDD-  234
                                *:  ****:**:.  * *:******::: *  ****** **:   .

U2AF2A_Fbor        ED--NPTAYLNGIVSTVVQDSPNKIFVGGLPNYLNEDQVKELLIAFGPLRAFNLVKDTTT  345
U2AF2B_Fbor        EDMALPQIHVKGIVSNFVKDSVNKIFLGGLPNYMNDEQVKELLEAFGPLRGFSLVKDSTT  342
U2AF2_Fpel         -DLGLPEIHLKDVVGTVVRDSVNKVFIGGLPSYLNDDQVKELLQAFGPLRSFNLVKDSTT  293
                    *   *  :::.:*...*:** **:*:****.*:*::****** ******.*.****:**

U2AF2A_Fbor        GLSKGYAFCEYVDAQITDQSIAGLHGMQLGEKKLIVQRAALGSKTGA-AMTAPVTIQVPG  404
U2AF2B_Fbor        GFSKGFGFAEYVDTGVTDMAIAALHGMEISERRLVVQRAELGVKGDPLSKHGPTPIQVPG  402
U2AF2_Fpel         GFSKGFGFCEYVDVSVTDQAIAGLHGMMLGERRLVVQRAAVGQKGDPMAKHGPTSIQVPG  353
                   *:***:.*.****. :** :**.**** :.*::*:**** :* * .  :  .*. *****

U2AF2A_Fbor        AQQALQNMKDTRPTEVLCLLNMVTVEELAEDEEYEDIVADIKEECEKFGEVKSIEIPRPT  464
U2AF2B_Fbor        LNMT-VATKESLPTRVVCLINVVTLEELRDEEDYEDILDDMKDECGKYGRVKSIEIPRPV  461
U2AF2_Fpel         INFT-EAIKESKPTTVVCLLNMVTQEELKDDEEYEDIVEDIHEECGKYGAVKSIEVPRPM  412
                    : :    *:: ** *:**:*:** *** ::*:****: *:::** *:* *****:***

U2AF2A_Fbor        QGMADLGGLGKIYVEFGEIGQCMECSNALAGRKFSQRVVMTMYYDPDKYHRRVFE519
U2AF2B_Fbor        AGE-EVGGLGKVYVEFAAVGDSIKATNSLSGRKFAQRVVMTCYYDEERFNLRDFE515
U2AF2_Fpel         HGV-EVGGLGKIYVEFEQTDDCIKANNALAGRKFSQRVVMTMYYDPDKFHRRMFE466
                    *  ::*****:****   .:.::..*:*:****:****** *** :::: * **
```

**Figure S2:** Multiple sequence alignment of *F. borealis* U2AF2 paralogues with the *F. pellucida* U2AF2. Color highlight shows strongest similarity between *F. pellucida* U2AF2 and *F. borealis* U2AF2α (yellow) or U2AF2β (green).

```
                                                                  Zinc Finger 1
U2AF1β_Fritillaria_borealis          -MHNGSGAHHLSSIFGTERDKVNCTFYYRIGTCRYGDRCPKVHNRPTFSPTIMIKNMYMN     59
U2AF1α_Fritillaria_borealis          --MDGSGAEYLASIFGTEKDKVNCSFYYKMGACRHGDRCSRVHNRPTFSQTILIQNFYQN     58
U2AF1_Fritillaria_haplostoma         --MDGSGAEYLASIFGTEKDKVNCSFYYKIGACRHGDRCSRVHNRPTFSQTILIQNLYKN     58
U2AF1_Oikopleura_dioica              -MDADAGAEYLASIFGTEKDKVNCSFYFKIGACHYGDRCSRLHNKPTFSQTMLLQNLYRS     59
U2AF1_Oikopleura_vanhoeffni          MTDADAGAEYLASIFGTEKDKVNCSFYFKIGACHYGDRCSRLHNKPTFSQTMLLKNLYRS     60
U2AF1_Strongylocentrotus_purpuratus  ------MAEYLASIFGTEKDKVNCSFYFKIGACRHGDRCSRLHNKPTFSQTIVIQNIYHN     54
U2AF1_Saccoglossus_kowalevskii       ------MAEYLASIFGTEKDKVNCSFYFKIGACRHGDRCSRLHNKPTFSQTIVLLNVYQN     54
U2AF1_Phallusia_mammillata           ------MAEYLASIFGTEKDKVNCSFYFKIGACRHGERCSRLHNKPTFSQTILIQGLYQN     54
U2AF1_Ciona_intestinalis             ------MAEYLASIFGTEKDKVNCSFYFKIGACRHGERCSRLHNKPTFSQTIVLQGLYQN     54
U2AF1_Ciona_savignyi                 ------MAEYLASIFGTEKDKVNCSFYFKIGACRHGDRCSRLHNKPTFSQTIVLQGLYQN     54
U2AF1_Branchiostoma_belcheri         ------MAEYLASIFGTEKDKVNCSFYFKIGACRHGDRCSRLHNKPTFSQTILMPNLYRN     54
U2AF1_Branchiostoma_floridae         ------MAEYLASIFGTEKDKVNCSFYFKIGACRHGDRCSRLHNKPTFSQTILMPNLYRN     54
U2AF1_Human                          ------MAEYLASIFGTEKDKVNCSFYFKIGACRHGDRCSRLHNKPTFSQTIALLNIYRN     54
U2AF1_Danio_rerio                    ------MAEYLASIFGTEKDKVNCSFYFKIGACRHGDRCSRLHNKPTFSQTIALLNIYRN     54
                                            *.:*:******:*****:**:::*:*::*:** ::**:**** *: : ..* .
                                                       RRM
U2AF1β_Fritillaria_borealis          PKLSSFNTSGEV-------CNMNEAEMQEHFDAFFDEIYTELEDSYGEIDEMIICDNLGE     112
U2AF1α_Fritillaria_borealis          PALSGLGSAGEN-------CDLGEIEMQEHYDGFFDEVYSELEEKYGEIEEMNVCDNLGD     111
U2AF1_Fritillaria_haplostoma         PALTNFNKDGEI-------CEMSELEMQEHYDSMFLEIYEEIERKYGEVEEMNICDNLGD     111
U2AF1_Oikopleura_dioica              PILDDDSG-----------QQITETMLQQHYDEFYEEVFFEIEDKYGEIEEMNVCENLGD     108
U2AF1_Oikopleura_vanhoeffni          PILDDNGG-----------MQITESMLQKHYDEFYEEVFFEIEDKYGEIEEMNVCENLGD     109
U2AF1_Strongylocentrotus_purpuratus  PANTAQSAEGGAAVGIADQSNLSDVEVQQHFDDFFEEVFAEMEQKYGEIEEMNVCDNLGD     114
U2AF1_Saccoglossus_kowalevskii       PQNSAQTAEGTH------FKGMSDVDMQKHFDDFFEEVFTELDDKYGEIEEMNVCDNIGD     108
U2AF1_Phallusia_mammillata           PQINNT-----A------AGTFTDVEMQEHYDNFFEEVFVELEEKYGSIEEMNVCDNLGD     103
U2AF1_Ciona_intestinalis             PQINNT-----G------IGTVSDVEMQEHYDNFFEEVFVELEEKYGSIEEMNVCDNLGD     103
U2AF1_Ciona_savignyi                 PQINNT-----G------MGTVSDVEMQEHYDNFFEEVFVELEEKYGSIEEMNVCDNLGD     103
U2AF1_Branchiostoma_belcheri         PQNSAQTADGSH------LSNISDVEMQEHYDNFFEEVFTELEEKYGEIEEMNVCDNLGD     108
U2AF1_Branchiostoma_floridae         PQNSAQTADGSH------LSNISDVEMQEHYDNFFEEVFTELEEKYGEIEEMNVCDNLGD     108
U2AF1_Human                          PQNSSQSADGLR-------CAVSDVEMQEHYDEFFEEVFTEMEEKYGEVEEMNVCDNLGD     107
U2AF1_Danio_rerio                    PQNTAQSADGLN--------AVSDVEMQEHYDEFFEEVFTEMEEKYGEVEEMNVCDNLGD     106
                                     *                    . :  :*:*:* :: *:: *:: .**.::** :*:*:*:
                                                           UHM                Zinc Finger 2
U2AF1β_Fritillaria_borealis          HLCGNVFCMFKDEDDAFKCVKALNDRWVDGRPIQAELCPITDFRDACCRQYEFGDCNRGG     172
U2AF1α_Fritillaria_borealis          HLVGNVYVKFKDEEMSQKAVKEINNRWFDKKPIYGELSPVSDFREACCRQYELGECNRGG     171
U2AF1_Fritillaria_haplostoma         HLVGNVYIKFKDEDESMKAVKELNERWFDKFPVYSELSPVSDFREACCRQYELGECNRGG     171
U2AF1_Oikopleura_dioica              HLVGNVYIKFYREEDCEKAAEGVNDRWFGMQPIVGELSPVTNFREACCRDYEMGECTRGG     168
U2AF1_Oikopleura_vanhoeffni          HIVGNVYIKFYKEEDCEKTAEMCNERWFAGQPVEAELSPVTNFREACCRDYEMGECTRGG     169
U2AF1_Strongylocentrotus_purpuratus  HLVGNVYVKFRYEEDAEKAVEDLNNRWFNRQPIRAELSPVTDFREACCRQYEMGECTRGG     174
U2AF1_Saccoglossus_kowalevskii       HLVGNVYVKFRNEEDAEKAVNDLNNRWFNGQPIRAELSPVTDFREACCRQYEMGECTRGG     168
U2AF1_Phallusia_mammillata           HLVGNVYVKFHKEEDAEKAVNDLNNRWFNGQPILSELSTVTDFREACCRQYEMGECTRGG     163
U2AF1_Ciona_intestinalis             HLVGNVYIKFHKEEDAEKAVNDLNNRWFNGQPIMSELSTVTDFREACCRQYEMGECTRGG     163
U2AF1_Ciona_savignyi                 HLVGNVYIKFHKEEDAEKAVNDVNNRWFNGQPIMSELSTVTDFREACCRQYEMGECTRGG     163
U2AF1_Branchiostoma_belcheri         HLVGNVYVKFRYEEDAEKAVKDLNNRWFNGQPIHAELSPVTDFREACCRQYEMGECTRGG     168
U2AF1_Branchiostoma_floridae         HLVGNVYVKFRYEEDAEKAVKDLNNRWFNGQPIHAELSPVTDFREACCRQYEMGECTRGG     168
U2AF1_Human                          HLVGNVYVKFRREEDAEKAVIDLNNRWFNGQPIHAELSPVTDFREACCRQYEMGECTRGG     167
U2AF1_Danio_rerio                    HLVGNVYVKFRREEDAEKAVINLNNRWFNGQPIHAELSPVTDFREACCRQYEMGECTRGG     166
                                     *: ***:  *  *: . * .   *:**.   *: .**. :::**:****:**:*:*.***
                                     Zinc Finger 2
U2AF1β_Fritillaria_borealis          LCNFMHIKPISREMRKELHRRYKRKKREDKKFRDDTASGDKGGYAGGFGGWSGIGDDYWD     232
U2AF1α_Fritillaria_borealis          FCNFMHLKPISRQLRKALYKRNKHKKSEDKSVRKHTAGGRHGGYAGGFGGWSGVGDGYND     231
U2AF1_Fritillaria_haplostoma         FCNFMHLKPISRELRKYLYKKKKKRHEDKQFREETAGGRHGGYAGGFGGWAGVGDGYDD     231
U2AF1_Oikopleura_dioica              FCNFMHLKPISKDLRKKLYAKKDKHRGYNP-DRRRGGRSRS--RSP-------------     211
U2AF1_Oikopleura_vanhoeffni          FCNFMHLKPISKALRKKLYNRKE-RRGYNP-DRRRGGRSKS--RSN-------------     211
U2AF1_Strongylocentrotus_purpuratus  FCNFMHLKPISRELRRELYGRRKRKKRSSRSKSTSPGPRDN--KRR-------------     218
U2AF1_Saccoglossus_kowalevskii       FCNFMHLKPISRELRRELYGRRVRTKK-RSRSRDRKSRSRS--RER-------------     211
U2AF1_Phallusia_mammillata           FCNFMHLKPISRELRRQLYGRKEKKRKSRS------PPTRS--K-R-------------     200
U2AF1_Ciona_intestinalis             FCNFMHLKPISRELRRRLYGRKEKKKRSRS------PPTRN--R-R-------------     200
U2AF1_Ciona_savignyi                 FCNFMHLKPISRELRRRLYGRKEKKRRSRS------PPTRN--R-R-------------     200
U2AF1_Branchiostoma_belcheri         FCNFMHLKPISRELRRELYGRRRRRRRSRSRDRSRSPPTRG--SRR-------------     212
U2AF1_Branchiostoma_floridae         FCNFMHLKPISRELRRELYGRRRRRRRSRSRERSRSPPTRG--SRR-------------     212
U2AF1_Human                          FCNFMHLKPISRELRRELYGRRRKKHRSRSRSRERRSRSR--------------------     207
U2AF1_Danio_rerio                    FCNFMHLKPISRELRRELYGRRRKRHRSRSRSRERRSRSR--------------------     206
                                     :*****:****: :*: *: :     :

U2AF1β_Fritillaria_borealis          RDHGFLGRQSTQSASKRKLDRIDNLLYGGGN-QGGG-----NQGGNHGGNHDRNQGGGQS     286
U2AF1α_Fritillaria_borealis          QDYGYDGRSSTQYAAERKFDKMEADKSKIGGHRGYQHHGHDQQGYGQQGHEQQGYGGQNG     291
U2AF1_Fritillaria_haplostoma         RDYGYYGKQSTQFGADRKLKKIDRDMKDKGGREGGGGNHHG--GGGNRQDNGGNYGGNRG     289
U2AF1_Oikopleura_dioica              ----------------------GRRRDR----DG-------------------------     219
U2AF1_Oikopleura_vanhoeffni          ----------------------SPRRRD----GGGG-----GGGGGGGRHGG---GGGG     237
U2AF1_Strongylocentrotus_purpuratus  ----------------------SRSRSR----SRGK-----------------------     228
U2AF1_Saccoglossus_kowalevskii       ----------------------SRRSRS----RSRDKRRSRSR-----------------     228
U2AF1_Phallusia_mammillata           ----------------------SRSRE-----RHRRH----------------------     210
U2AF1_Ciona_intestinalis             ----------------------GGSRE-----RHGRDRRH-------------------     213
U2AF1_Ciona_savignyi                 ----------------------GSKER-----HRSRERR--------------------     212
U2AF1_Branchiostoma_belcheri         ----------------------SRSHE-----RGGGSRRSRSRERGGRGKAGG---GG--     240
U2AF1_Branchiostoma_floridae         ----------------------SRSHE-----RGGGSRRSRSRERGGRGKAGG---GGGG     242
U2AF1_Human                          ------------------------DRG-----RGGGGG-----------------GGGG     220
U2AF1_Danio_rerio                    ------------------------ERN-----RGGGGGG----GGGGGGGGG---GGGG     230

U2AF1β_Fritillaria_borealis          RG-----AHRNVDDRSRERAASQRPR--------     307
U2AF1α_Fritillaria_borealis          QRGYYPEH--------------------------     299
U2AF1_Fritillaria_haplostoma         GGGNYKREYRGEGDAEYGGNK--RPREMNFR---     318
U2AF1_Oikopleura_dioica              ------------R-NGRGGRGRRHQPDLDWSLNY     240
U2AF1_Oikopleura_vanhoeffni          GGGRYG--GRDRG-PPRGGRGGRHQPDLDWSLNY     268
U2AF1_Strongylocentrotus_purpuratus  -------DDRHRE-RERSGRF-------------     241
U2AF1_Saccoglossus_kowalevskii       ---SKDKDHRSRS-RERSGRF-------------     245
U2AF1_Phallusia_mammillata           ----------------------------------     210
U2AF1_Ciona_intestinalis             ----------------------------------     213
U2AF1_Ciona_savignyi                 ----------------------------------     212
U2AF1_Branchiostoma_belcheri         ---GSRSRSRSRS-RERSGRF-------------     257
U2AF1_Branchiostoma_floridae         GGAGSRSRSRSRS-RERSGRF-------------     262
U2AF1_Human                          GGGRERDRRRSRD-RERSGRF-------------     240
U2AF1_Danio_rerio                    GGGRDRERRRSRD-RERSGRF-------------     250
```

| | |
|---|---|
| RRM | RNA recognition motif |
| UHM | U2AF homology motif |
| (yellow) | U2AF1β-specific mutations in Zinc fingers |

**<u>Figure S3:</u>** Multiple sequence alignment of *F. borealis* U2AF1 paralogues with deuterostome U2AF1.

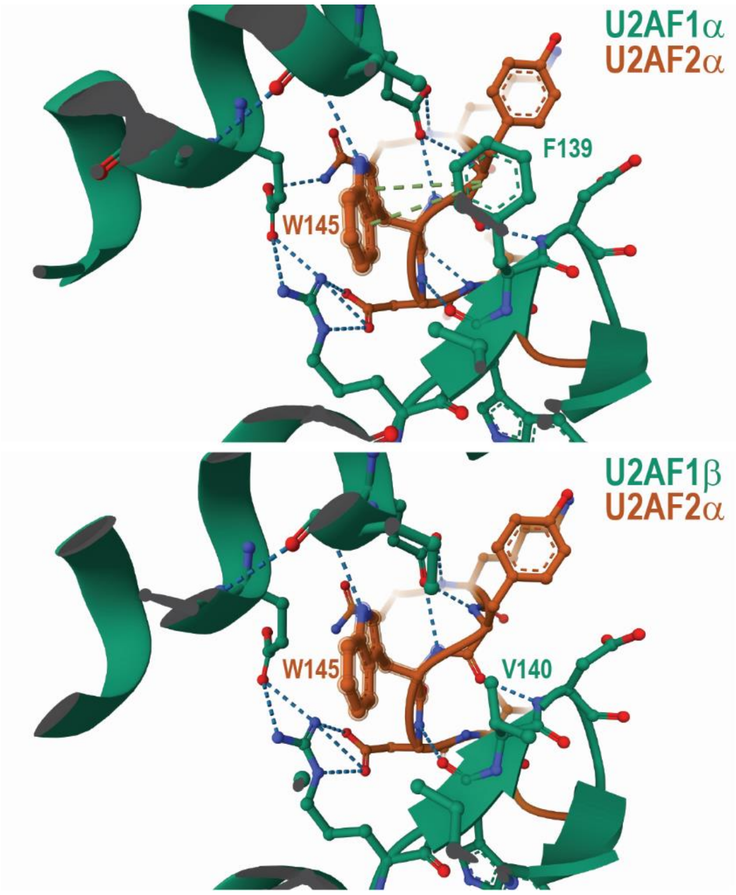


**<u>Figure S4:</u>** Models of complexes formed between *F. borealis* U2AF2α and U2AF1α (top) or U2AF1β (bottom), showing the UHM/ULM interface. Interactions between aromatic rings are shown with dashed green lines, dashed blue lines represent hydrogen interactions. Structures were predicted with AlphaFold 3[1] and selected based on best alignment with the crystal structure of human U2AF[2].

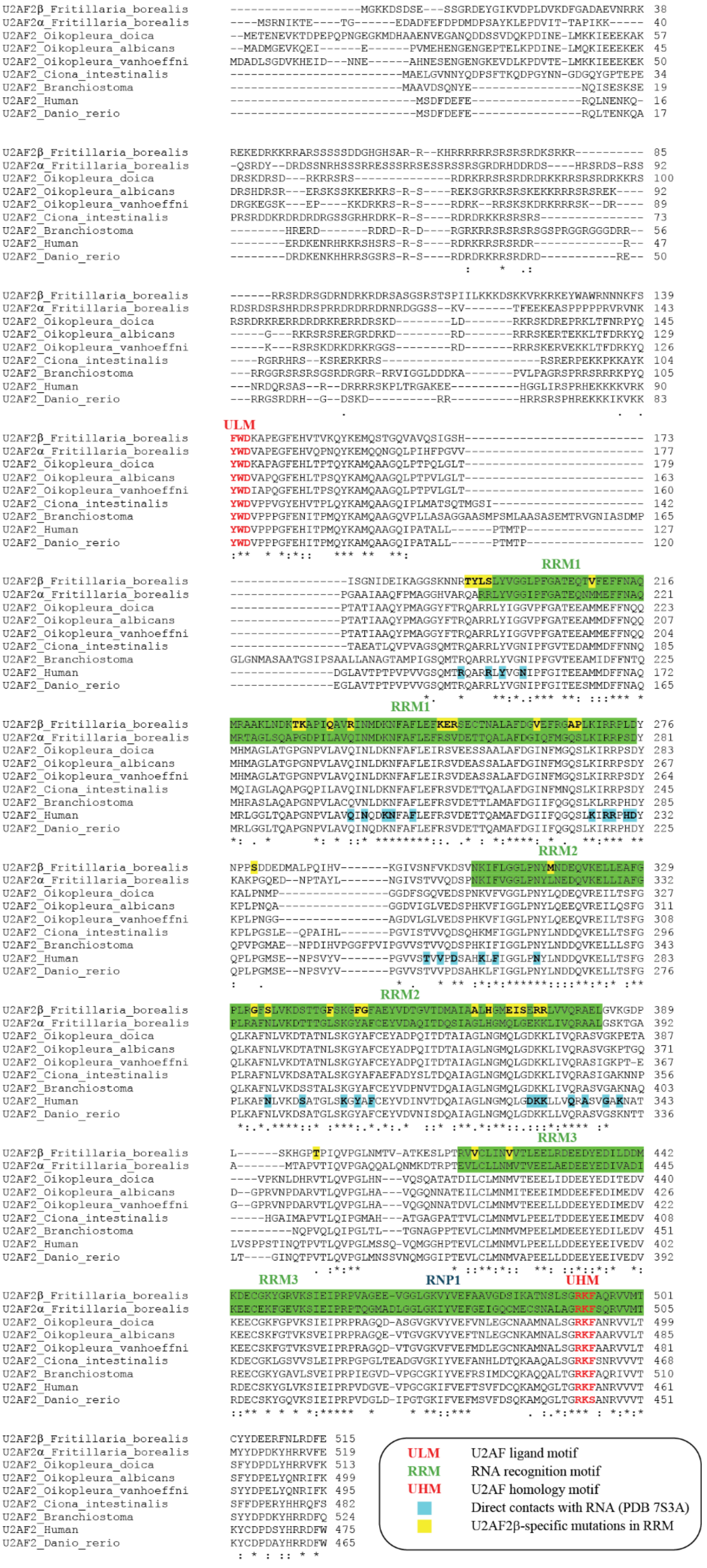
ULM
RRM1
RRM2
RRM3
RNP1
UHM
ULM U2AF ligand motif
RRM RNA recognition motif
UHM U2AF homology motif
Direct contacts with RNA (PDB 7S3A)
U2AF2β-specific mutations in RRM

**<u>Figure S5:</u>** Multiple sequence alignment of *F. borealis* U2AF2 paralogues with deuterostome U2AF2.

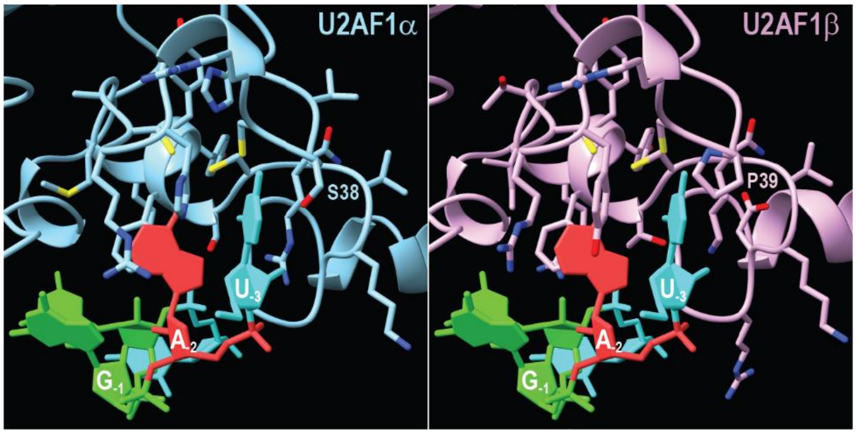


**<u>Figure S6:</u>** Models of *F. borealis* U2AF1 paralogues bound to UAGGU, showing how amino acid change in the first zinc finger could impact the recognition of 3'ss. The models were established with SWISS-MODEL[3], using the crystal structure of yeast U2AF bound to UAGGU as a template[4].

| $1^{st}$ base | $2^{nd}/3^{rd}$ base: AA | AC | AG | AU | AA | AC | AG | AU |
|---|---|---|---|---|---|---|---|---|
| A | 1,2 | 1,4 | 1,4 | 0,5 | 0,8 | 0,9 | 0,3 | 1,5 |
| C | 2,5 | 1,5 | 0,7 | 1,1 | 1,7 | 0,9 | 1,1 | 1,2 |
| G | 0,4 | 1,1 | 1 | 0,6 | 0,6 | 0,9 | 1,3 | 1 |
| U | 0,7 | 1,1 | 0,7 | 0,6 | 1,1 | 1,1 | 1,8 | 1,5 |
| | U2AF1β vs U2AF1Od | | | | U2AF1Od vs U2AF1β | | | |

| $1^{st}$ base | $2^{nd}$ base: A | C | G | U | A | C | G | U |
|---|---|---|---|---|---|---|---|---|
| A | 1,1 | 1,3 | 0,9 | 0,7 | 0,9 | 0,9 | 1,1 | 1,4 |
| C | 1,3 | 1,7 | 1,3 | 1,3 | 1 | 1,3 | 0,8 | 1,2 |
| G | 0,9 | 1,5 | 0,4 | 0,9 | 1,1 | 0,8 | 0,9 | 1,1 |
| U | 0,8 | 1,2 | 0,9 | 0,7 | 1,3 | 1,3 | 1,1 | 1,2 |
| | U2AF1β vs U2AF1Od | | | | U2AF1Od vs U2AF1β | | | |

**Figure S7:** Results obtained with *F. borealis* U2AF1β and *O. dioica* U2AF1 during RNAcompete experiments. High affinity oligos were identified by comparing RNA enriched in one sample but depleted in the other sample. The tables show the abundance of 3- and 2-mer motifs in the sequence of high affinity oligos. Values correspond to the average frequency in collected oligos relative to the average frequency measured in the initial library.

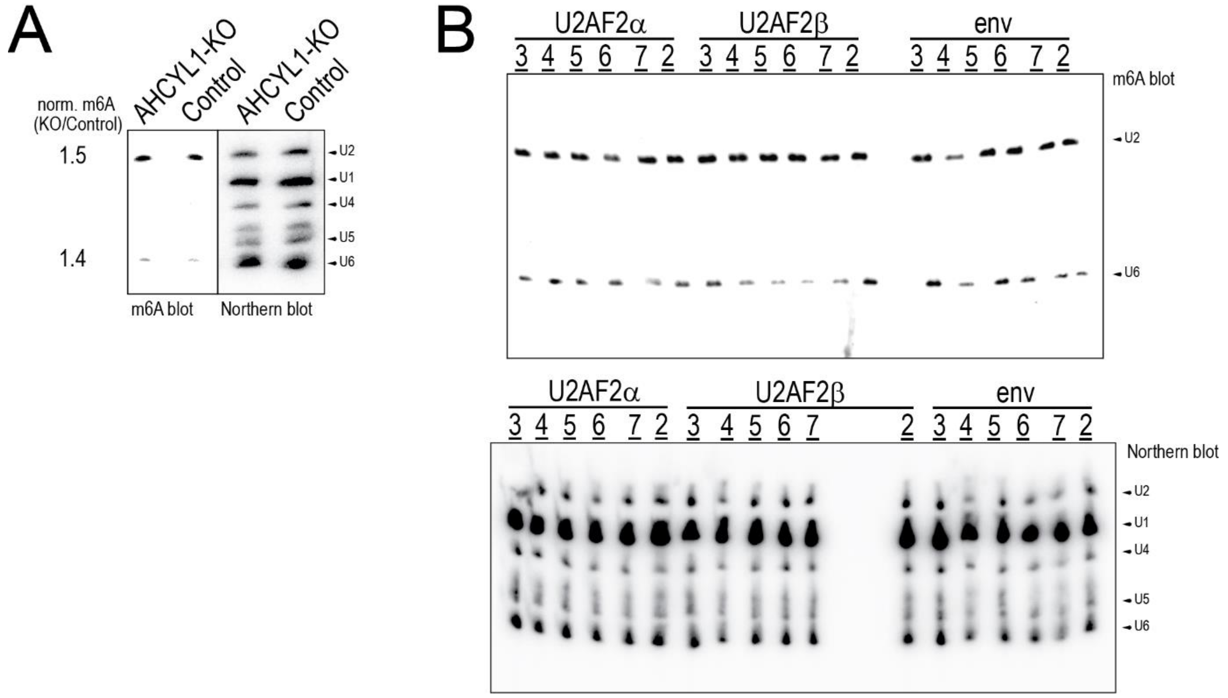


**<u>Figure S8:</u>** Experimental data used to measure the effect of AHCYL1 and U2AF2 on m6A modification of snRNA. **A)** m6A immunoblot and snRNA Northern blot of HEK293T cells engineered with a Crispr-cas9 KO of AHCYL1, compared to control cells. To measure m6A levels, the m6A signals corresponding to either U2 or u6 snRNA were normalized against the snRNA expression measured with northern blot. **B)** m6A immunoblot and snRNA Northern blot of HEK293T cells transfected with *F. borealis* U2AF2 paralogues, or an Env control. We established a series of six replicates (lanes labelled 2-7) for each transfection. For each replicate, m6A levels were measured as described in **A)**.

| | *F. borealis 2018* | *FLYE* | *FLYE-MERGE* | *FLYE-FINAL* |
|---|---|---|---|---|
| Source | Single individual | Pooled individuals | Pooled individuals | Pooled individuals |
| Approach | Illumina | ONT | ONT + Illumina | ONT + Illumina |
| Size (bp) | 89.645.637 | 77.586.534 | 79.706.325 | 79.691.002 |
| Contigs | 32.542 | 4.060 | 4.065 | 4.063 |
| Avg contig size (bp) | 2754 | 19.109 | 19.617 | 19.613 |
| Largest contig (bp) | 138.349 | 251.966 | 251.770 | 251.709 |
| N50 (bp) | 10.252 | 38.885 | 39.913 | 39.913 |
| Ns | 5.318.489 | 0 | 4.474 | 4.474 |
| Gaps | 10.613 | 0 | 15 | 15 |

**Table S1:** Statistics and approaches to establish *F. borealis* genome assemblies. Column 1 corresponds to the initial Illumina-based assembly[5], column 2 corresponds to an assembly based on ONT sequencing, column 3 corresponds to a hybrid merging Illumina and ONT assemblies, column 4 corresponds to the final assembly used to produce the data in this study.

| Termini | A-U | A-C | G-G | G-U | A-A | U-U | A-G | G-C | C-U | C-C | U-G | U-A | C-A | U-C | G-A | C-G |
|---|---|---|---|---|---|---|---|---|---|---|---|---|---|---|---|---|
| tWW | 3.71 | **2.47** | **0.00** | **3.14** | **1.09** | **3.41** | *n.o.* | 6.49 | 5.40 | 4.00 | **3.24** | 3.77 | **2.58** | 5.42 | *n.o.* | 6.54 |
| cWH | **3.42** | *n.o.* | **0.00** | *n.o.* | *n.o.* | **2.58** | **2.85** | *n.o.* | 3.67 | **2.76** | 7.25 | 5.64 | *n.o.* | *n.o.* | **2.22** | 5.41 |
| Frequency | 0.23 | 0.22 | 0.10 | 0.07 | 0.06 | 0.05 | 0.04 | 0.03 | 0.03 | 0.03 | 0.03 | 0.03 | 0.02 | 0.02 | 0.02 | 0.02 |

**Table S2:** Base pairing between intron termini of *F. borealis*. First and second rows indicate, for the tWW and the cWH geometries, the isodiscrepancy between each pair of intron termini and the G-G reference base pair. Third row shows the frequency of intron termini in the genome.
